# Architecture and evolutionary conservation of *Xenopus tropicalis* osteoblast-specific regulatory regions shed light on bone diseases and early skeletal evolution

**DOI:** 10.1101/2023.08.30.555543

**Authors:** Héctor Castillo, Francisco Godoy, Clément Gilbert, Felipe Aguilera, Salvatore Spicuglia, Sylvain Marcellini

## Abstract

Understanding the genetic mechanisms underpinning the differentiation of osteoblasts, the bone producing cells, has far reaching implications for skeletal diseases and evolution. To this end, it is crucial to characterize osteoblastic regulatory landscape in a diverse array of distantly-related vertebrate species. By comparing of the ATAC-seq profile of *Xenopus tropicalis* (*Xt*) osteoblasts to liver, heart and lung control tissues, we identified 524 promoters and 6,750 distal regions whose chromatin is specifically open in osteoblasts. Nucleotide composition, Gene Ontology, and RNA-Seq confirmed that the identified elements correspond to *bona fide* osteogenic transcriptional enhancers, and TFBS enrichment revealed a well-conserved regulatory logic with mammals. Amongst the 357 *Xt* osteoblast-specific enhancers aligning to homologous human loci, 127 map to regions annotated as enhancers. Phenotype predictions based on the genes neighbouring these conserved enhancers are tightly related to impaired skeletal development. In addition, six conserved enhancers are located at loci associated to craniosynostosis (*mx2*, *tcf12*), osteopoikilosis (*lemd3*), osteopenia (*gorab*), skeletal dysplasia (*flnb*) and craniofacial abnormalities (*gpc4*). From an evolutionary perspective, the elephant shark genome aligns to 53 *Xt* osteoblast-specific enhancers that are also conserved and annotated as enhancers in humans, revealing an ancestral osteogenic role for the ATOH8, IRX3, NFAT, NFIB and MEF2C transcription factors, as well as for the FGF, IHH and BMP/TGFb signalling pathways. As the absence of bone in sharks is a derived feature, we propose that, in this lineage, the osteogenic regulatory network has been maintained for its function in odontoblasts. Our data argues in favour of a common origin for dentine and bone, and provides a glimpse into the key regulatory elements and upstream activators that drove the formation of an ancient type of mineralized tissue in the vertebrates that inhabited the oceans more than 460 million years ago.

**Author Summary:** During animal embryogenesis, distinct type of tissues are formed and assembled, resulting in an integrated, functional organism. During this process, cells must make important decisions, which largely rely on an accurate use of their genetic material. Here, we have studied how the genome “knows” that it must participate to the formation of the bone tissue in a frog animal model. We therefore identified important genomic regions that are involved in driving the expression of genes involved in the formation of a mineralized skeleton. On the one hand, we show that some of these regions are also present in humans, and, therefore, skeletal pathologies could be studied in the frog model at a genetic level. On the other hand, we also identify regions that are present in the genome of a shark, which allows us to propose an evolutionary framework for the early evolutionary origin of the vertebrate skeleton.

## Introduction

One daunting challenge of the post-genomic era is to bridge the gap between genotype and phenotype to better understand developmental processes, evolutionary trajectories, and the etiology of complex diseases [1, 2]. Cracking open the genome’s regulatory code involves understanding how distinct cell lineages acquire their specific characteristics through the progressive activation of regulatory enhancers and transcriptional programs [3, 4]. In this respect, a major effort has been made in identifying and classifying regulatory elements at a genome-wide scale. For instance, more than a decade ago, the advent of ChIP-Seq technology allowed a series of pioneering chromatin profiling experiments to be performed in flies and mammals [5–8]. More recently, the ATAC-Seq technology has provided straightforward access to the repertoire of open chromatin regions genome-wide, which are often associated with regulatory potential [9]. Hence, ATAC-Seq has been used to generate open chromatin atlases of diverse mammalian tissues [10–18]. The fact that a fraction of distant enhancers is rapidly evolving, and cannot readily be identified by sequence similarity searches, emphasizes the importance of generating regulatory element databases in a wide variety of species [16].

With its sequenced and assembled genome, the amphibian *Xenopus tropicalis* (*Xt*) represents a powerful model organism to improve our understanding of human genetic disorders [19–25]. Recently, ATAC-Seq has been performed in this species to explore gene regulatory networks involved in early development [26, 27] and tail regeneration [28]. In *Xt*, osteogenesis follows a conserved pattern with mammals, as mesenchymal precursors progressively differentiate into osteoblasts, the bone producing cells [29, 30]. However, amphibians have never been employed to examine the osteoblastic regulatory landscape, which has so far only been studied in great depth in mammals [31–35]. In mouse, a combination of genome wide approaches, knock-out animals, and reporter experiments, has revealed how signalling pathways and transcription factors cooperate at enhancers to drive ossification in a well-controlled spatio-temporal fashion [35–46]. In parallel, GWAS studies have allowed the identification of polymorphism associated with skeletal diseases [47–49]. Identifying bone-specific regulatory elements and upstream activators in *Xenopus tropicalis*, and, more particularly, those components that are conserved with humans, would represent a valuable resource for improving our understanding of bone development and disease in this amphibian organism which is readily amenable to transgenesis and site-directed mutagenesis [19, 21, 23, 24].

In addition to its biomedical relevance, the bone tissue is particularly fascinating from an evolutionary perspective, as it represents a major innovation of the vertebrate group by fulfilling diverse functions related to body protection and mineral reserves [50]. Indeed, detailed analyses of the dermal skeleton from fossils from the Silurian-Devonian period (443-360 Mya) have shown that bone emerged long before the radiation of the jawed vertebrates whose extant representative include both the chondrichtyans (*e.g.* elephant shark, sharks and rays) and the osteichthyans (*e.g.* coelacanth, lungfish, gar, teleost fish, amphibian and amniote groups) [50, 51]. One fruitful strategy to explore the early origin of the osteoblastic cell type has been to compare the expression of genes coding for bone matrix components between jawed vertebrate representatives [30, 52–55]. While the evolutionary origin of the osteoblastic gene regulatory network has attracted much attention in the literature [56–58], it still lacks formal experimental assessment, and the nature of the ancestral osteogenic regulatory elements and of their upstream input remains unknown. Solving this issue is instrumental to understand the origin of the osteoblast, and, more broadly, to shed light on the genetic mechanisms involved in the evolution of new animal cell types [59].

Here, we compared the ATAC-Seq profile of *Xenopus tropicalis* native osteoblastic cells to a variety of non-mineralizing control tissues such as larval heart, liver and lung. Taken together, our data allows us to (i) describe the architecture of the *Xt* osteoblast-specific promoters and enhancers, (ii) identify distal enhancers conserved with human genes involved in skeletal diseases, and (iii) probe the evolutionary origin of the regulatory network involved in the emergence of the vertebrate mineralized skeleton.

## Results

### Identification of osteoblast, liver, heart and lung open chromatin regions in *Xenopus tropicalis*

To characterize the nucleosome-free regions (NFRs) of *Xt* osteoblastic cells, we worked with freshly extracted, uncultured, primary osteoblasts obtained following a previously established protocol [30]. Our analysis also included larval liver, heart and lung as soft tissues that served as useful controls to discriminate between ubiquitous and osteoblast-specific regulatory regions. The osteoblastic ATAC-Seq experiments showed the expected pattern of fragment size distribution, with discrete peaks corresponding to the NFR, monucleosome and dinucleosome fractions (Fig 1A). By focusing on annotated *Xt* transcription start sites (TSSs), we found that the ATAC-Seq NFR and monucleosome signals form two distinct clusters. The first cluster is composed of 5,949 promoters harbouring a robust NFR located immediately upstream of the TSS and flanked by two well-positioned nucleosomes (Fig 1B, left panel), whereas the second cluster contains 16,947 promoters showing weak NFR and diffuse mononucleosome signals (Fig 1B, right panel). Similar situations were observed for the liver, heart and lung tissues (S1 Data). After performing peak calling using the NFR signal, peaks for the four tissues were merged into a standard peak list, the number of raw reads mapped to each standard peak was counted, and Pearson’s correlations were calculated, depicting a high correlation between the two replicates of each sample (Fig 1C). We next examined the distribution of peaks with respect to gene bodies (Fig 2). When considering all the peaks from each tissue (Fig 2A), we found a relatively consistent pattern, with 26-39% of peaks located at TSSs, 26-34% of intronic peaks, 24-29% of intergenic peaks, and a minority of peaks falling into exons and transcription termination sites (TTSs). Calculating the distance of all the non-TSS peaks from each tissue to their closest TSS also revealed a consistent distribution pattern (Fig 2B). The osteoblastic non-TSS NFRs were located slightly more distally than the other samples, with 32% of the peaks included within 5kb of the closest annotated promoter, and 53% of the peaks located further away than 10kb of the closest annotated promoter (Fig 2B).

**Figure 1:**
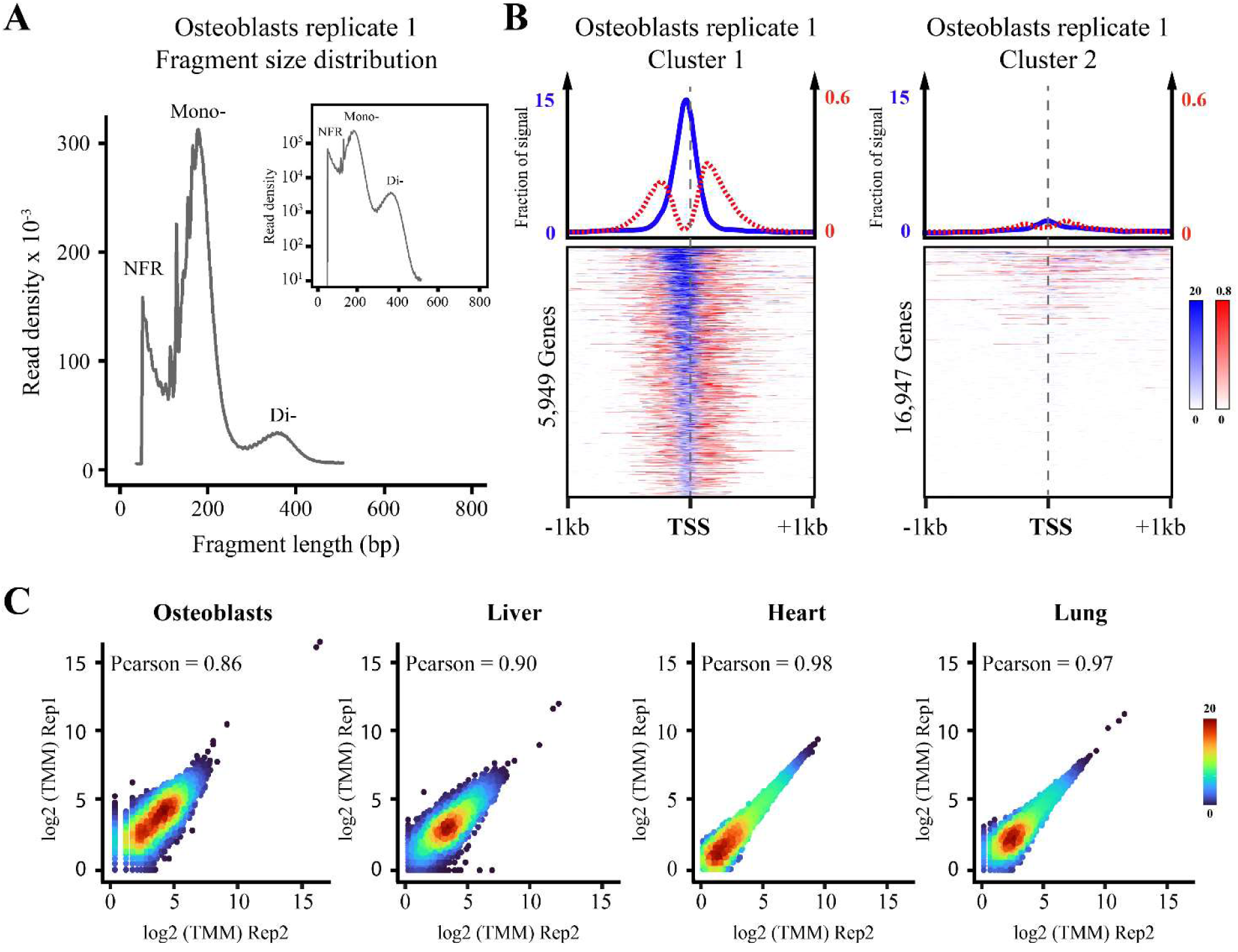
Characterization and validation of the ATAC-Seq libraries. **(A)** Fragment size distribution in linear and logarithmic (inset) scales for the first replica of the bone sample. **(B)** The NFR (blue) and mononucleosome (red) signals at annotated *Xt* TSSs are separately visualized and clustered, revealing a cluster of 6,342 TSSs harbouring a clear NRF flanked by two well-positioned nucleosomes (left) and a cluster of 18,249 TSSs lacking a NFR and displaying a diffuse mononucleosome signal (right). **(C)** Correlation of the NFR signal for both replicas from each of the examined tissues. Pearson correlation values are shown.

**Figure 2:**
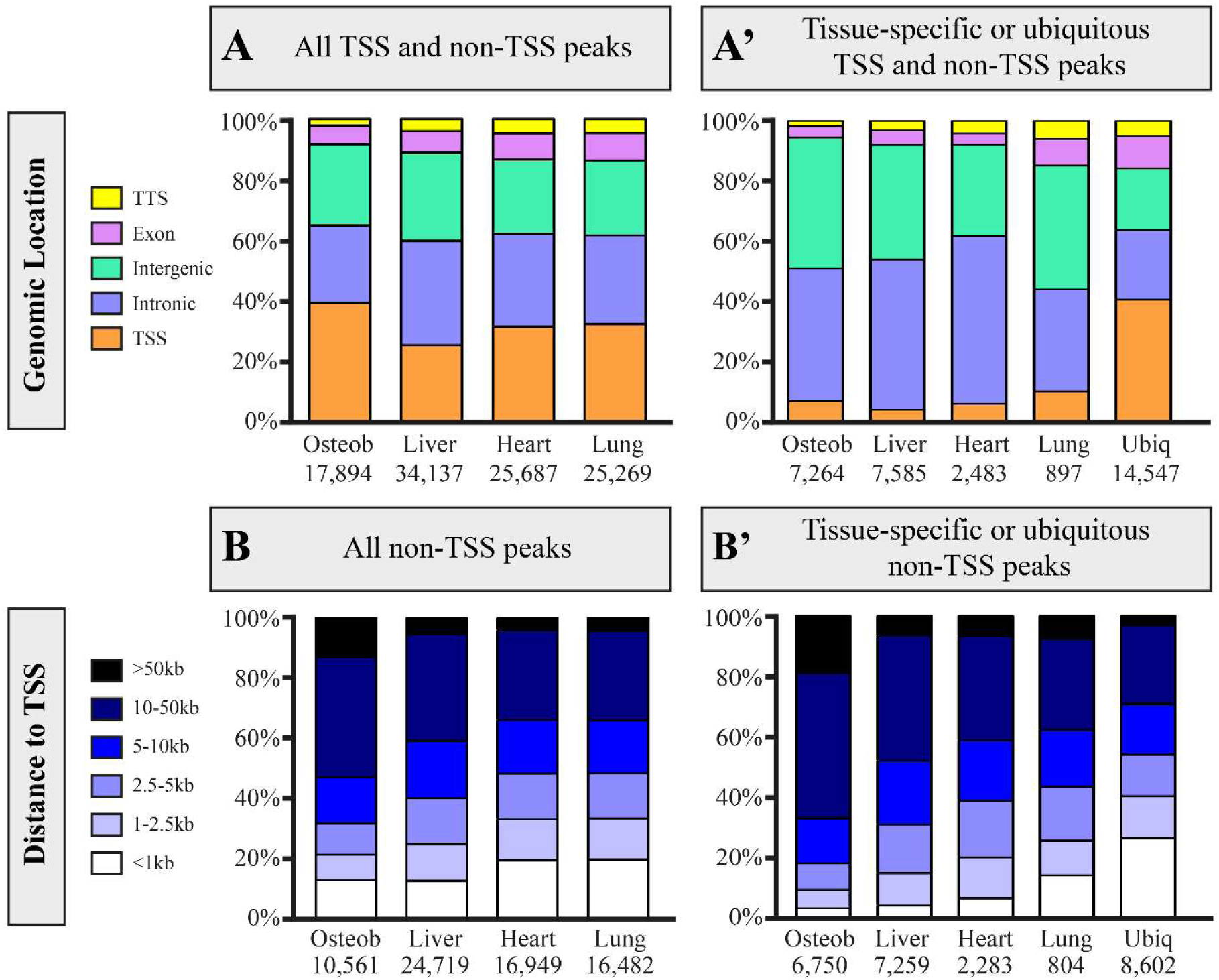
Spatial distribution of the identified ATAC-Seq peaks with respect to annotated gene bodies in *Xenopus tropicalis.* Top: percentage of peaks falling in the indicated types of annotated genomic regions, either considering all the peaks identified for each tissue **(A)** or only considering tissue-specific or ubiquitous peaks **(Á)**. Bottom: percentage of non-TSS peaks located at the indicated distance from the closest annotated TSS, either considering all the non-TSS peaks identified for each tissue **(B)** or only considering tissue-specific or ubiquitous non-TSS peaks **(B’)**. In each case, the number of open chromatin regions identified by peak calling is indicated. Abbreviations: TSS, transcription start site; TTS, transcription termination site; Osteob, osteoblasts; Ubiq, ubiquitous.

Globally, the Pearson correlation coefficient was much higher for TSS than non-TSS peaks (Compare Fig 3A and B). This finding is consistent with previous studies showing that, between distinct cell types, histone marks are largely invariable at promoters while they display highly context-dependent patterns at enhancers [6, 7]. Hierarchical clustering showed that the landscape of the NFRs from heart and lung are more similar to each other than to osteoblasts or liver, which is true both for TSS and non-TSS regions (Fig 3A and B). By contrast, the global distribution of the osteoblastic NFRs was the most distinct from the any of the three other assayed tissues (Fig 3A and B).

**Figure 3:**
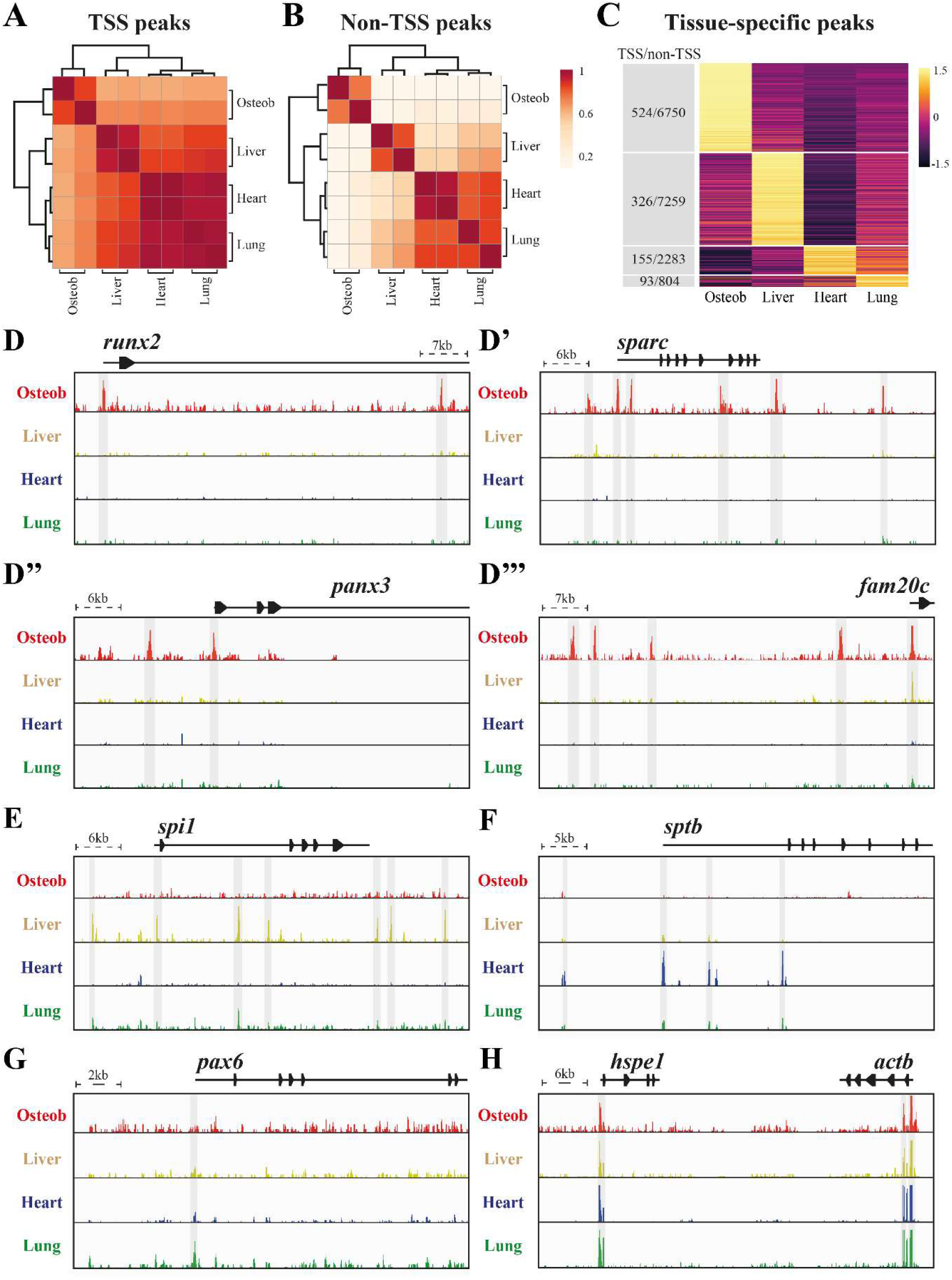
Identification of tissue-specific TSS and non-TSS open chromatin regions in *Xenopus tropicalis*. Heatmap showing the Pearson correlation coefficient and hierarchical clustering of all the ATAC-Seq peaks identified in a given sample and annotated as TSS **(A)** and as non-TSS **(B)** in the *Xt* genome. **(C)** Clustering analysis of ATAC-Seq peak signal intensity uncovers tissue-specific peaks. For each tissue, the number of TSS and non-TSS open chromatin regions is indicated on the left. **(D-H)** Illustrative snapshots of genomic loci exhibiting ATAC-Seq peaks specific to bone (D-D’’’), liver (E), heart (F), lung (G), or shared among all tissues (H). For each locus, the scale, gene names, transcription units (horizontal lines), and coding exons (triangles) are indicated. Grey shades indicate open chromatin regions showing a significant enrichment by peak calling.

### Characterization of tissue-specific and ubiquitous nucleosome-free regions in *Xenopus tropicalis*

We reasoned that subtracting the osteoblastic NFRs with data from a variety of non-mineralized control tissues, such as liver, heart and lung, would lead to a substantial enrichment in specific NFRs closely related to the activation of genes involved in osteogenesis and bone matrix deposition. To this end, we clustered of all the peaks from the four samples and identified ubiquitous and tissue-specific and TSS and non-TSS peaks (Fig 3C, S2 Data). To illustrate that our procedure identifies osteoblast-specific peaks, figure 3D-D’’’ shows snapshots of the NFR signal at loci of genes known to be involved in bone development and evolution such as *runx2*, *sparc*, *panx3* and *fam20c* [60–64]. Our approach also reveals the presence of tissue-specific peaks for the liver (Fig3E), heart (Fig 3F) and lung (Fig 3G), as well as ubiquitous peaks that display NFRs in all tissues (Fig 3H).

Figure 2Á shows that, with respect to their genomic locations, tissue-specific peaks are mainly located in introns (34-55%) and intergenic regions (30-43%), while ubiquitous peaks are predominantly found at promoters (41%). When considering only the non-TSS regions, tissue-specific peaks are located much more distally than ubiquitous peaks (Fig 2B’). We found a strong correlation between the NFR and mononucleosome signals around tissue-specific and ubiquitous TSS and non-TSS peaks, meaning that open chromatin regions are consistently flanked by two well-positioned lateral nucleosomes (S3 Data).

Gene Ontology analyses on genes whose promoters are most-closely located to the identified non-TSS peaks revealed biological processes and molecular functions that are highly consistent with each category of tissue-specific or ubiquitous peaks (Fig 4). Hence, osteoblast-specific regions (Fig 4A) are enriched in skeletogenic genes involved in collagen matrix deposition, osteoblast differentiation, as well as TGFbeta and integrin-mediated signaling pathways that are known to positively control osteogenesis [65–67]. By contrast, liver-specific regions are associated with genes related to the immune system and macrophage biology, while heart-specific regions are associated with erythrocytes and blood vessels and, to a lesser extent, with fibroblasts and cardioblasts (Fig 4B and C). These results suggest that our tissue-dissociation procedure enriched the liver and heart samples in white and red blood cells, respectively, and is consistent with the fact that the larval liver is an important site of myelopoiesis in *Xenopus* [68]. Finally, ubiquitous non-TSS peaks are located close to genes involved in fundamental cell biology processes, including transcription, cell cycle, chromatin organization and vesicle transport (Fig 4D). Due to the low number of regions, no significant results were obtained with lung-specific non-TSS ATAC-Seq peaks, nor with any category of TSS (data not shown).

**Figure 4:**
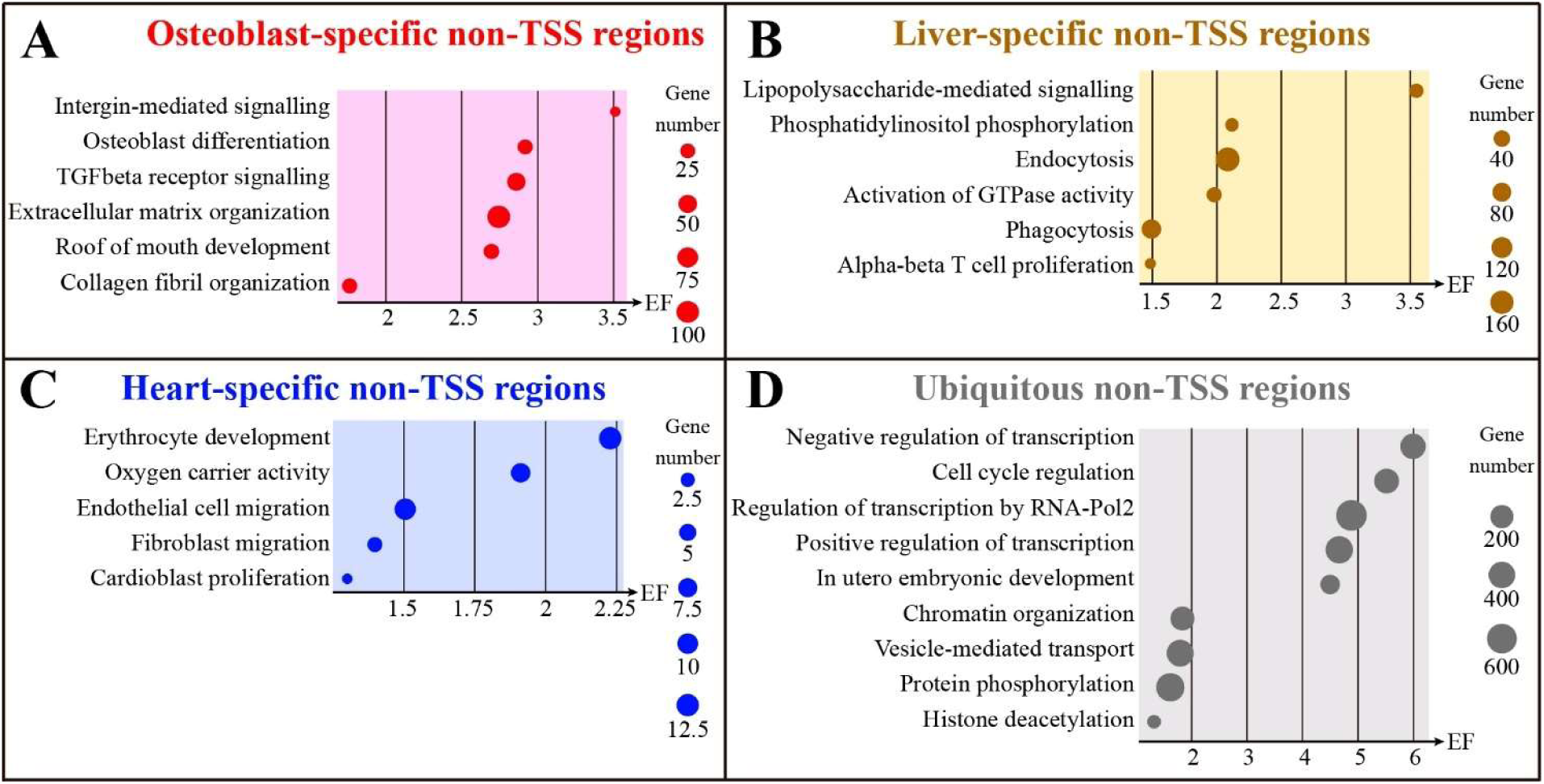
Gene ontology analyses of genes associated with non-TSS regions. Gene ontology terms are shown for osteoblast-specific (**A**), liver-specific (**B**), heart-specific (**C**) and ubiquitous (**D**) non-TSS open chromatin regions. Putative target genes whose promoter is located the closest to the identified ATAC-Seq peaks were selected for GO analyses. For each term, the graphs show the number of associated genes (circle size) and the enrichment factor (EF) in the x-axis.

We conclude that our comparative ATAC-Seq experiment allowed the accurate identification of distal non-TSS NFR regions that display key features expected for transcriptional enhancers because they are found in the vicinity of osteogenic genes and are located much more distally than ubiquitous non-TSS peaks.

### Sequence features of tissue-specific and ubiquitous open chromatin regions in *Xenopus tropicalis*

To further understand the nature of the identified open chromatin regions, we focused our analysis on their underlying sequence features. We first examined the CpG content because this dinucleotide has been associated with regulatory regions [69–71]. We found that all TSS and non-TSS regions display a central CpG enrichment (Fig 5A). In particular, ubiquitous and lung-specific regions display the strongest CpG enrichment in TSS (around 5% CpG) and non-TSS (around 2.5% CpG) regions (Fig 5A). CpG enrichment is the weakest for liver-specific TSS regions and for heart-specific and liver-specific non-TSS regions, where the peak in CpG frequency just reached random levels and was largely due to lateral depletion in this dinucleotide (asterisks in Fig 5A). We separately examined the frequency in the C and G nucleotides and found that the enrichment is always higher at TSS than at non-TSS regions (Fig 5B). In addition, our analysis revealed a higher C and G nucleotide frequency for ubiquitous and lung specific TSS (around 30%) and non-TSS (around 26%) regions (Fig 5B). A strong GC-skew phenomenon is evident only for ubiquitous TSS (and, to a lesser extent, also for the lung tissue, see arrowheads in Fig 5B), which is consistent with the association of GC-skewing with non-methylated, highly expressed, promoters [72]. Finally, a lateral depletion in C and G frequency is also observed for the osteoblast-specific, heart-specific and liver-specific non-TSS regions (asterisks in Fig 5B). While the reason for lateral depletion and central enrichment of C, G and CpG is unclear, we note that the binding of the CEBPA transcription factor is related to a similar CpG pattern in the macaque and dog genomes [73].

**Figure 5:**
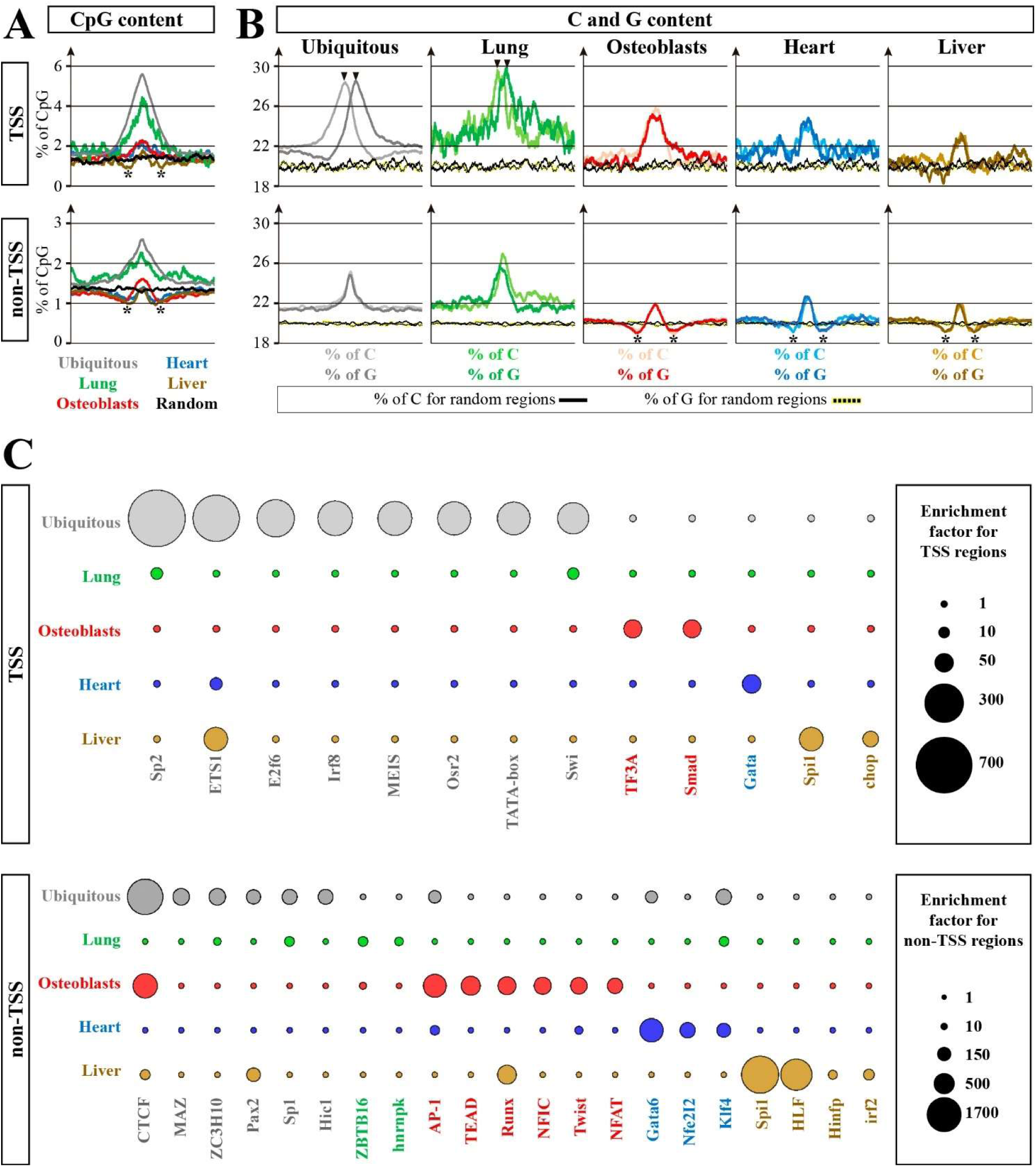
Underlying sequence features of TSS and non-TSS elements. Percentage of the CpG dinucleotide **(A)** and of the C and G nucleotides **(B)** in 2kb regions centred on ATAC-Seq peaks at TSS (top) and non-TSS (bottom) regions. Results are shown separately for ubiquitous (grey), lung-specific (green), bone-specific (red), heart-specific (blue), and liver-specific (brown) open chromatin regions, as well as for randomly selected control sequences (black). All TSS were orientated with the transcription units pointing rightward. Black arrowheads indicate a higher proportion of C and a higher proportion of G upstream and downstream of the TSS, respectively (GC-skew). Asterisks indicate a depletion in frequencies that drop below control levels on each side of the peak. **(C)** Enrichment in TFBSs as TSS (top) and non-TSS (bottom) regions. The circle size represents the enrichment factor as indicated on the right. A colour code is associated with the transcription factor names in function of the sample where their TFBS is most highly enriched.

We then analyzed each type of TSS and non-TSS region for transcription factor binding site (TFBS) enrichment (Fig 5C and S4 Data). A detailed description of these findings is shown in S5 Data, and, for the sake of clarity, we will focus here on the most crucial aspects of TFBS enrichment found at ubiquitous and osteoblast-specific NFR. For instance, ETS, SP and E2F6 have been reported to function at ubiquitous and/or highly transcribed promoters [74–77], which is consistent with an enrichment for their TFBS in ubiquitous TSS in our dataset (Fig 5C, top panel). Interestingly, ubiquitous non-TSS regions are enriched in TFBSs for CTCF and MAZ which have been shown to bind and collaborate at insulators and chromatin borders [78, 79]. By contrast, osteoblast-specific TSS regions are enriched for SMAD TFBS, in agreement with an important role of the BMP pathway in this cell type [38]. Finally, osteoblast-specific non-TSS regions are enriched in TFBS of many well-known osteogenic regulators in mammals (Fig 5C, bottom panel, S5 Data), including the AP-1 member FRA1 [39], TEAD [40], RUNX2 [35], NFIC [41], TWIST1/2 [42] and NFAT [43].

Taken together, the analyzed sequence features support the idea that we accurately identified each type of TSS and non-TSS regulatory region. Hence, we expect ubiquitous non-TSS regions to contain a mixture of ubiquitous enhancers and insulators, while the osteoblast-specific non-TSS dataset might be highly enriched in osteogenic enhancers.

### The expression strength of genes associated with tissue-specific open chromatin regions is context-dependent

To obtain evidence that the identified regions contain active promoters and enhancers involved in transcriptional activation, we assessed whether the NFR presence correlates with gene expression in osteoblasts and larval liver. We chose these two samples because they contain the highest number of identified regions of interest (Fig 3C) and display the most specific NFR signal (S3 Data). In addition, they differ in their CpG and C/G profiles (Fig 5A and B) and are enriched in distinct sets of TFBSs (Fig 5C, and S4, S5 Data), which makes the comparison informative from a sequence perspective. Hence, RNA-Seq was performed for osteoblasts and larval liver, and gene expression profiles were compared between the two cellular contexts (Fig 6). For TSS (Fig 6A, lines 1-6), we exclusively used genes whose promoter overlaps with a tissue-specific or ubiquitous peak. We found that genes associated with osteoblast-specific TSS peaks are expressed more robustly in osteoblasts than in the liver (Fig 6A, lines 1 and 2). Conversely, genes associated with liver-specific TSS peaks are expressed more strongly in the liver than in osteoblasts (Fig 6A, lines 3 and 4). Finally, genes associated with ubiquitously opened TSS are expressed at similar levels in both tissues (Fig 6A, lines 5 and 6). For non-TSS regions (Fig 6A, lines 7-12), we exclusively used genes whose promoter was located closest to a tissue-specific or ubiquitous peak. Although the nearest gene is not necessarily the target of the identified enhancer, we found that, like for TSS, the non-TSS peaks display a strong tissue-dependent behavior. Hence, genes associated with osteoblast-specific non-TSS peaks are expressed more strongly in osteoblasts than in the liver (Fig 6A, lines 7 and 8). The opposite is true for liver-specific non-TSS peaks (Fig 6A, lines 9 and 10). Finally, only marginal expression difference (*p* = 0.026) was observed, between each sample, for genes whose promoter is located close to a ubiquitous non-TSS peak (Fig 6A, lines 11 and 12). Taken together, these data strongly support the notion that we successfully identified TSS and non-TSS NFR associated with transcriptionally active genes, and, most importantly, that the set of osteoblast-specific non-TSS NFR corresponds to *bona fide* osteoblastic enhancers.

**Figure 6:**
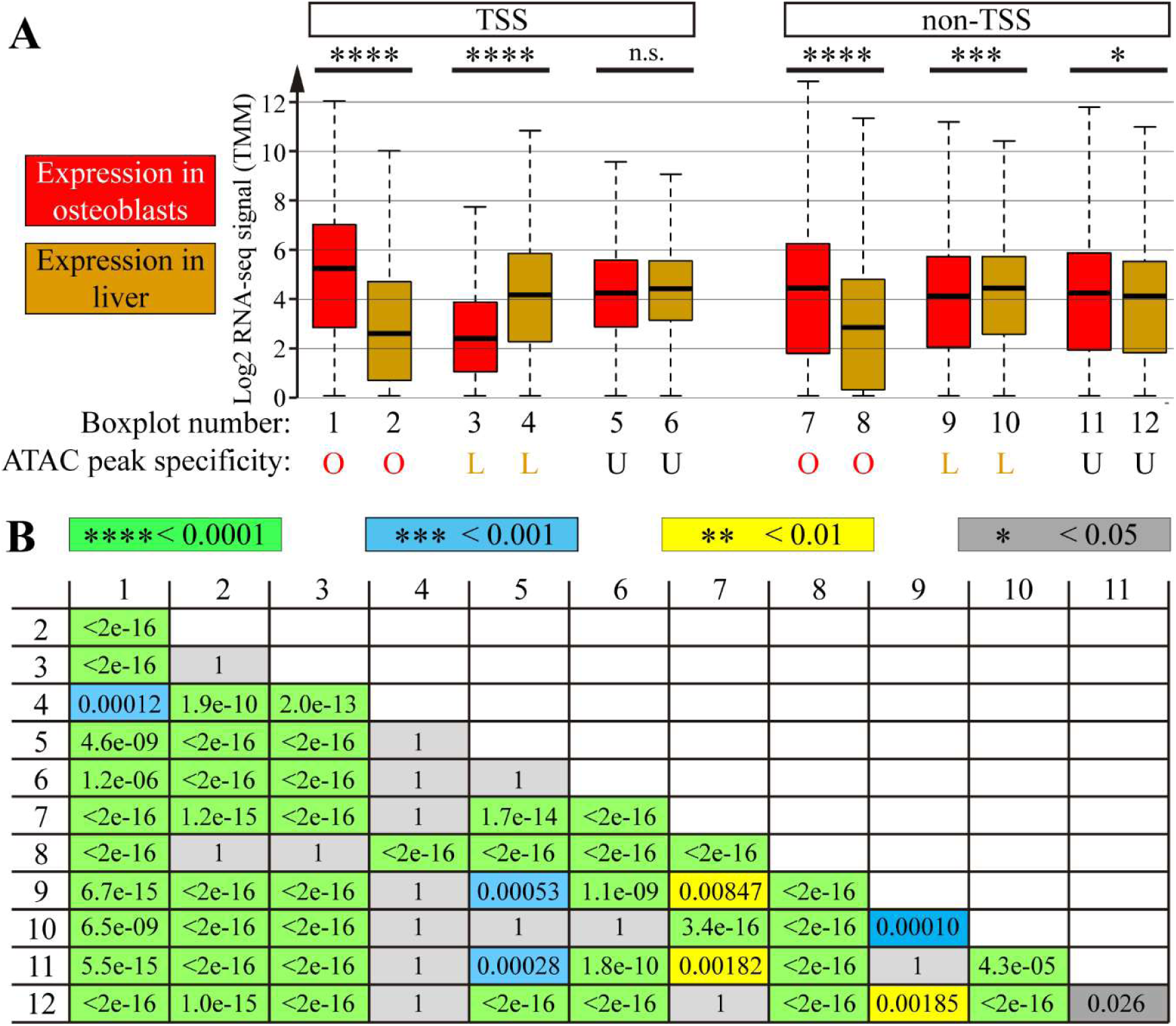
Expression values of genes associated with ATAC-Seq peaks. **(A)** The boxplots show the distribution of RNA-Seq TMM values (Log2) for genes expressed in osteoblasts (red) or liver (brown). We show the expression for genes that are associated with an ATAC-Seq peak at the level of their TSS (boxplots 1-6) or genes that are most closely located to a non-TSS ATAC-Seq peak (boxplots 7-12). ATAC-Seq peaks were either osteoblast-specific (O, boxplots 1, 2, 7, and 8), liver-specific (L, boxplots 3, 4, 9, and 10), or ubiquitous for the four tissues examined in this study (U, boxplots 5, 6, 11 and 12). Statistical results are indicated for the same type of ATAC-peaks in distinct cellular contexts (n.s., non-significant). Outliers were omitted for the sake of clarity. **(B)** Statistical analyses of the distribution of expression values between all 12 boxplots. A non-parametric Kruskal Wallis test and a Bonferroni P correction were used.

### Conservation degree of *X. tropicalis* tissue-specific and ubiquitous open chromatin regions among chordates

To explore the biomedical and evolutionary relevance of our dataset, we assessed the degree of conservation of the identified NFRs with humans as well as with six other species belonging to the chordate phylum (Fig 7 and S6 Data). For this purpose, we used *Xt* NFRs as queries to perform a series of similarity searches using BLASTN against the genomes of the clawed frog *Xenopus laevis* [a pipidae frog which diverged from Xt 48 Mya, 80], of two amniote species [human and chick, which diverged from Xt around 351-333 Mya, 81], of the zebrafish *Danio rerio* [a teleost fish which diverged from Xt around 458-449 Mya, 81], of the elephant shark *Callorhinchus milii* [a chondrichthyan showing mineralized skeletal elements, which diverged from Xt shortly before the split with the teleost fish lineage, 61, 81, 82, 83], of the lamprey *Petromyzon marinus* [a cyclostome, devoid of any type of mineralized tissues, which diverged from Xt around 500 Mya, 50, 84], and of the amphioxus *Branchiostoma floridae* [a non-vertebrate cephalochordate species which diverged from Xt around 550 Mya, 50, 85]. For the sake of comparison, we also performed BLASTN analyses using as queries the lung-specific, heart-specific, liver-specific and ubiquitous regulatory elements, and also conducted the same analysis with two sets of randomly selected regions of the *Xt* genome whose length matches the average length of the NFR at TSS and non-TSS (S6 Data). We computed an enrichment factor defined as the percentage of hits obtained with any type of NFR divided by the percentage of hits obtained with random regions. This analysis showed that, globally, the osteoblast-specific and lung-specific NFR are well-conserved (with various enrichment factor superior to 2), the liver-specific NFR are particularly divergent (with various enrichment factor inferior to 0.5), and the heart and ubiquitous NFR exhibit an intermediate conservation level, often yielding an enrichment factor close to 1 (Fig 7). Taken together, this data is consistent with the notion that the conservation degree of enhancer elements can vary greatly depending on their tissue-specificity [86].

**Figure 7:**
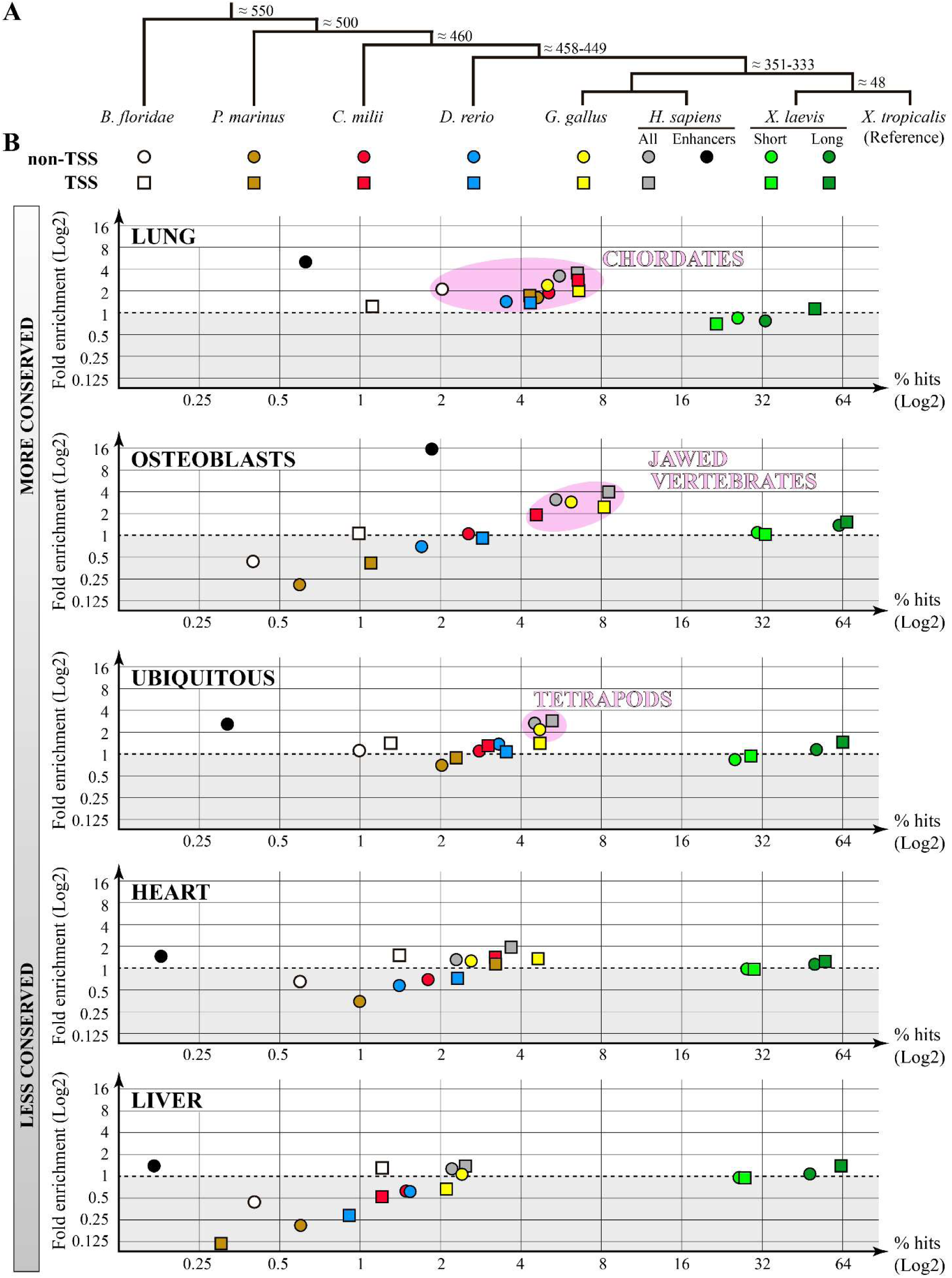
Sequence conservation of *Xt* open chromatin regions among seven chordate species. **(A)** A cladogram shows the phylogenetic relationships of the amphioxus *Branchiostoma floridae*, the lamprey *Petromyzon marinus*, the elephant shark *Callorhinchus milii*, the zebrafish *Danio rerio*, the chicken *Gallus gallus*, the human *Homo sapiens*, the clawed frog *Xenopus laevis* and *Xenopus tropicalis*. Estimated divergence time (Millions of years) is indicated for each node. **(B)** The conservation degree of *Xt* non-TSS (circles) or TSS (squares) ATAC-Seq peaks were evaluated by BLASTN against the genomes of the aforementioned species. When blasting *H.s.* with non-TSS *Xt* peaks, we either considered the total number of hits (“all”, grey), or only the subset of hits aligning to human regions annotated as an enhancer (black). When blasting the allotetraploid frog *X.l.*, we considered separately the short (light green) and the long (dark green) subgenomes. For each category of *Xt* ATAC-Seq peaks (*i.e.* tissue-specific or ubiquitous), the graphs show the percentage of regions that were alignable by BLASTN (x-axis), as well as the enrichment factor compared to a selection of random *Xt* regions (y-axis). For ease of comparison, the area where enrichment values are inferior to 1 (dotted line) is shaded in light grey. The pink ellipses highlight clusters of conserved non-TSS and TSS regions, and the indicated name refers to the phylogenetic group at the base of which at least a fraction of these regulatory elements can be inferred to have emerged.

### Identification osteoblast-specific regulatory regions relevant for understanding human osteogenesis and skeletal pathologies

Of a total of 524 osteoblast-specific TSS peaks, 48 align to the promoter of their corresponding homologous gene in the human genome (Fig 7, S6 and S7 Data), representing a 4.8 fold increase with respect to random. By performing a literature search, we found that 19 of these genes have been associated to the osteogenic and skeletogenic processes (S7 Data), including genes coding for critical osteogenic transcription factors such as RUNX2 and SIX2 [64, 87], and for important bone matrix components such as ADAM12 and TYPE III COLLAGEN [88, 89].

Amongst the 6,779 osteoblast-specific non-TSS peaks, 370 align to the human genome and, among these, 127 match to human regions annotated as enhancers, representing a 16-fold enrichment factor with respect to control (Fig 7, S6 and S8 Data). We found that synteny was conserved between *Xt* and human for 357 NFR (*i.e.* in 96.5% of the cases, S8 Data). None of the 13 osteoblast-specific non-TSS peaks that did not show conserved synteny were annotated as enhancers in humans, suggesting that they either represent false positives from our experiment, or that they act as long-range enhancers. We compared TFBS enrichment profile between the set of osteoblast-specific non-TSS regions that align to the human genome, and those that lack conservation, and found that binding sites for the previously identified osteogenic transcription factors (AP-1, NFAT, NFIC, RUNX, TWIST, TEAD, see Fig 5 and Data S4, S5) were enriched in both groups (S9 Data). One marked difference, however, was observed for the CTCF binding site that appears to be highly enriched only in non-conserved NFR (S9 Data). Taken together, this data suggests that the conserved regions correspond almost exclusively to transcriptional enhancers, while the non-conserved regions contain a combination of enhancers and insulators.

By applying Great on the 370 osteoblast-specific non-TSS NFR that are conserved between *Xt* and human, we obtained a strong association for human phenotypes related to anomalous skeletal development (S10 Data). Finally, all the genes located at the loci of these conserved regions were searched in a database of skeletal diseases [49], leading to the identification of putative osteogenic enhancers for *flnb*, *gorab*, *gpc4*, *lemd3*, *msx2* and *tcf12* (S11 Data). Snapshots of ATAC-Seq and RNA-Seq profiles at the *lemd3* (associated to the Buschke-Ollendorff syndrome), *msx2* (associated to Craniosynostosis 2), *tcf12* (associated to Craniosynostosis 3) and *gpc4* (associated to Keipert syndrome) loci are shown in Fig 8A.

**Figure 8:**
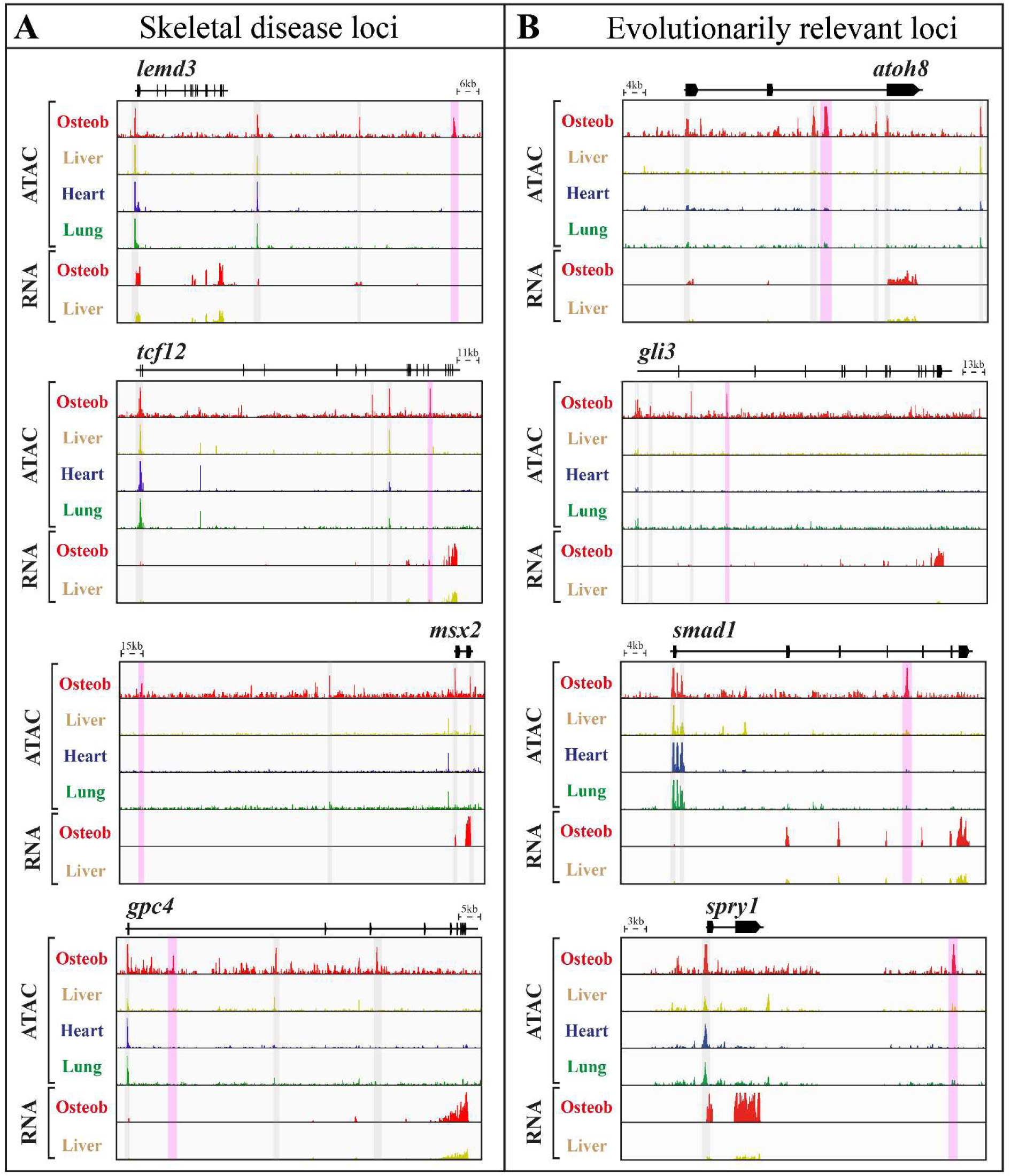
Conserved osteoblast-specific putative enhancers in genes of biomedical and evolutionary relevance. **(A)** Snapshots of *Xt* loci whose homologous human genes have been associated to skeletal diseases. Osteoblast-specific *Xt* non-TSS NFR that are conserved with the human genome are shaded pink. **(B)** Snapshots of *Xt* loci harbouring osteoblast-specific non-TSS NFR conserved with the human, chick and elephant shark genome, and that are also annotated as enhancers in humans (Pink shades). In (A) and (B), non-conserved osteoblast-specific NFR are shaded grey. See text for details and references.

### Comparison of the conserved *X. troplicalis* osteoblast-specific NFR with jawed vertebrates

Blasting the *Xt* osteoblast-specific TSS against the elephant shark genome recovered twice as many hits than expected by chance, leading to the identification of 15 promoters that are conserved between *Xt*, the elephant shark, chick and human (S12 Data). Of these, 7 promoters belong to genes associated with an osteogenic function in mammals, such as the emblematic *Runx2* gene [see S12 Data and Ref 64]. We identified 173 osteoblast-specific non-TSS NFR that align to the elephant shark genome, which only corresponds to an enrichment factor of 1.0 (S6 Data and Fig 7). Therefore, in an attempt to identify *bona fide* osteogenic enhancers that might have evolved at the base of the jawed vertebrates, we focussed our analysis on osteoblast-specific non-TSS sequences that are conserved between *Xt* and (i) elephant shark, (ii) chick, (iii) human, and (iv) that are also annotated as enhancers in the human genome. As shown in S13 Data, 53 open chromatin regions fulfil all these conditions, all of which displaying a conserved synteny between *Xt* and elephant shark. The location of these regions is biased towards transcription factors (18 genes, 12 of which having been reported to play a role during mammalian osteogenesis), and signalling pathway components (10 genes), including actors of the Fgf, Ihh, and BMP/TGFb signalling pathways which play instrumental roles during mammalian osteogenesis (S13 Data). Examples of ancient bone-specific enhancers located at the *atoh8*, *gli3*, *smad1* and *spry1* loci are shown in Fig 8B.

## Discussion

### On the nature of the open chromatin regions in non-mineralizing tissues

As native osteoblasts were obtained using a previously established extraction protocol [30], it is not surprising that this sample provides the most biologically consistent signal regarding Gene Ontology (Fig 4), TFBS enrichment (Fig 5, S4 and S5 Data), and the known osteogenic function of the mammalian genes associated with conserved TSS and non-TSS peaks (S7, S10, S11, S12 and S13 Data). By contrast, the cellular composition of the non-mineralizing tissues is less controlled, and GO and TFBS data suggest that the heart and liver mostly contain red and white blood cells, respectively, in agreement with the function of these two organs (Fig 4 and 5, and S5 Data). By far, the lung is the most puzzling tissue in this study as it exhibits unique characteristics that are not straightforward to interpret. This is because the lung-specific NFRs are very few (Fig 3), their nucleotide signature resembles that of ubiquitous elements (Fig 5), yet their patterns of conservation with other genomes are amongst the highest (Fig 7). In addition, the lung-specific non-TSS NFR are enriched in TFBS for HNRNP-K [a transcription factor involved in stem cell regulation and lung regeneration, see Refs 90, 91], and are located in the vicinity of genes known to play a role in stem cells of the lung and other tissues (S14 Data). We therefore propose that our dissociation protocol somehow led to a partial enrichment in lung pluripotent stem cells [92]. In conclusion, while the exact cellular composition of the three non-mineralizing control tissues remains unknown, the body of data provided here shows that they successfully served the purpose of identifying ubiquitous as well as osteoblast-specific TSS and non-TSS NFRs.

### Major differences in the architecture of tissue-specific and ubiquitous NFR in *Xenopus tropicalis*

We found that ubiquitous and tissue-specific NFR differ in many aspects, in agreement with data obtained in mammals. For instance, when compared to tissue-specific promoters, *Xt* ubiquitous promoters are clearly GC-skewed [72], they harbour a higher CpG content and control genes related to basic cellular functions [74, 93, 94], and they are enriched in TFBS for the ETS, SP and E2F6 transcription factors known to bind the TSS of broadly expressed genes [74–77]. In addition, only the ubiquitous non-TSS NFRs are enriched in both CTCF and MAZ TFBS, suggesting that many of them act as insulators [78, 79]. The osteoblast-specific non-TSS regions display some CTCF enrichment, which might correspond to tissue-specific insulators [95]. The fact that these CTCF sites are predominantly enriched in non-conserved regions (S9 Data) suggests that bone-specific insulators are evolutionarily more labile than enhancers, and, therefore, that osteoblast-specific chromatin domains might show substantial variation between frogs and humans. Finally, while the ubiquitous non-TSS NFR are mostly proximal, the distal location of *Xt* tissue-specific enhancers resembles the distribution of developmental enhancers [96], consistently with the idea that more complex regulatory architectures are required to generate specific expression patterns.

The presence of the TATA-box consensus in ubiquitous *Xt* promoters (Fig 5 and S4, S5 Data) clearly represents the most divergent feature with mammals where the presence of a TATA-box is associated with genes showing tissue-specific expression patterns [69]. The TATA-box being the only promoter motif conserved in all eukaryotic genomes examined to date, it is striking that it contributes to different promoter classes in amphibians and mammals. This shift in the use of the TATA-box motif might have been driven by the evolution of differences in nucleotide composition in these vertebrate lineages [97].

### Global conservation of regulatory regions

Our comparative analysis (Fig 7) revealed a series of noticeable features that are consistent with a body of published literature on other vertebrate species. First, we find that TSS regions are usually better conserved than non-TSS regions (as an example, human TSS regions always yield a higher percentage of hits and a higher enrichment factors than human non-TSS regions, compare the relative position of grey circles and grey squares on Fig 7), a situation that has also been observed between mammalian species having diverged 180 Mya [98]. Second, the identified *Xt* regulatory elements (as well as the random regions) align approximately twice as much to the *Xenopus laevis* L subgenome than to the S subgenome. This observation is consistent with the fact that the S subgenome is particularly prone to deletions and has significantly diverged [80]. Third, *Danio rerio* displays one of the most divergent genomes, often showing enrichment values inferior to those obtained when blasting the genomes of most remotely related species such as the elephant shark, lamprey, or even amphioxus. In this respect, it is worth mentioning that the genome of the gar (a non-teleost ray-finned fish) is much better conserved with tetrapods than the *Danio rerio* genome [99]. Fourth, some NFR display enrichment scores that are lower than expected by chance, a situation which is particularly evident when blasting the liver-specific NFR on the lamprey genome (Fig 8, bottom panel). If we consider that the liver sample led to the detection of *Xt* lymphocytic promoters and enhancers (Figs 4 and 5, S5 Data), the overall low conservation degree we observed might reflect a rapid divergence rate for the immune gene regulatory network since its origin at the base of the vertebrate more than 500 million years ago [100]. This idea is consistent with the absence of conservation for a macrophage enhancer at the *CSF1R* locus [101]. Fifth, the set of lung-specific enhancers aligns noticeably well to the amphioxus genome (2 fold enrichment score, see Fig 7 and S6 Data), considering that approximately 550 My separate this species from *Xt*. As amphioxus lacks lungs, this observation might reflect a conservation of enhancers related to stem cell function, as we have proposed above. This hypothesis supported by a study examining the regeneration process in amphioxus [102].

### A well-conserved regulatory logic for tetrapod osteoblastic enhancers

Remarkably, many of the TFBS enriched in the set of *Xt* osteoblast-specific NFR are bound by key osteogenic regulators in mammals such as SMAD1 at TSS, and AP-1, TEAD, Runx2, NFIC, TWIST1/2 and NFAT at non-TSS regions [35, 38–43], reflecting a highly conserved regulatory logic for osteogenesis at the tetrapod level. While the TWIST1 and 2 proteins have mainly been reported to delay the differentiation process by binding and inhibiting RUNX2 [42], one study showed that TWIST1 also directly repressed the *C-ROS-1* promoter, thereby negatively regulating osteogenic differentiation [103]. In this respect, the enrichment in TWIST1/2 TFBS at *Xt* osteogenic enhancers is a novel finding and might suggests that TWIST proteins control the onset of osteoblastic differentiation through redundant repressive mechanisms that act both at the post-translational and transcriptional levels. The only “new player” binding the osteoblast-specific promoters seems to be TFIIIA which controls the transcription of ribosomal genes associated with the Pol-III promoter in *Xenopus laevis* [104]. Hence, according to our work, TFIIIA might also regulate *Xt* Pol-II promoters during osteogenesis.

By examining 6,380 enhancers devoid of sequence conservation (S9 Data), we detected TFBS enrichment for the AP-1, NFAT, NFIC, RUNX, TWIST and TEAD transcription factors which are important regulators of osteogenesis in mammals (S5 and S9 Data). This shows that, during the 350 millions of years that separate amphibians from the amniote lineage, the osteoblastic enhancers have maintained a well-conserved regulatory grammar despite extensive sequence turnover, a phenomenon commonly observed in distinct animal groups and biological contexts [105–107]. This observation highlights the importance of experimentally characterizing osteoblastic distal enhancers in a variety of vertebrate species, as such regulatory elements cannot be exhaustively identified by sequence similarity searches.

### Identification of new osteogenic enhancers relevant for human biology

Here, we report 370 non-TSS osteoblast-specific NFRs that are conserved with humans (including 127 regions annotated as enhancers in the human genome), that are globally associated with a skeletogenic function (S8, S10 Data), and that are enriched in binding sites transcription factors known to play a major osteogenic role in mammals (S9 Data). By crossing our data with a recently updated nosology of genetic skeletal disorders [49], we identified 7 conserved enhancers at loci of genes involved in skeletal pathologies such as *lemd3*, *gpc4*, *tcf12*, *trps1*, *flnb*, *prrx1* and *msx2* (S11 Data, Fig 8A). Hence, the 370 conserved enhancers reported here could potentially be further studied in *Xt* to investigate the genetic bases of skeletal diseases.

### Evolutionary origin of the osteoblast-specific gene regulatory network

The *Xt* osteoblastic enhancers are particularly well-conserved (S6 Data, Fig 7), providing a unique opportunity to probe the evolutionary origin of the osteoblast-specific gene regulatory network. The location of the 53 *Xt* osteoblast-specific regions that are annotated as enhancers in humans and align to the human, chick and elephant shark genomes is biased towards genes encoding transcription factors and signalling pathway components (S13 Data). By focusing on genes with known osteoblastic roles in mammals, we propose that the ATOH8, BNC2, EBF3, FOXP1 and 2, HMGA2, IRX3, MEF2C, NFATC1, NFIB, ZEB2 and ZFHX4 transcription factors are ancient players involved in skeletal mineralization (S13 Data, Fig 8B). Likewise, enhancers located at the *spry1*, *gli3*, *ptch1*, *tgfbr3*, *smad1* and *bmp4* loci draw attention to the evolutionary importance of the Ffg, Hh and Bmp signalling pathways (S13 Data, Fig 8B). What could possibly be, however, the function of osteoblastic enhancers in a representative species of the cartilaginous fish, a group that lacks bone tissue and osteoblasts? This question is better answered if we first consider that the absence of bone is a secondary loss, and that cartilaginous fish evolved from a species that already possessed an ancestral form of bony tissue, similar to the plates covering the body of placoderms and osteostracan fossils [50, 51]. Second, it is important to note that bone and dentine share many similarities at the tissular, cellular and genetic levels [52, 61, 108, 109]. Here, we propose that osteoblasts and odontoblasts evolved from common a proto-skeletal cell that relied on the aforementioned transcription factors and signalling pathways. After the bone tissue loss in the chondrichthyan lineage, this ancestral regulatory network was maintained for its function in odontoblasts. According to this hypothesis, the genetic toolkit we uncovered might well have evolved long before the cartilaginous/bony fish split, around 460 million years ago, in the first cell type able to mineralize a rudimentary form of bone tissue.

## Materials and methods

### *Xenopus tropicalis* handling and bioethical considerations

The Ethics Committee of the University of Concepcion (Concepcion, Chile) approved all experimental procedures carried out during this study, which were performed following the guidelines outlined in the Biosafety and Bioethics Manual of the National Commission of Scientific and Technological Research (CONICYT, Chilean Government). *Xenopus tropicalis* tadpoles were staged according to the Nieuwkoop and Faber developmental table [110] and anesthetized with Tricaine (MS-222, Sigma), as recommended [111, 112].

### Sample collection and ATAC-Seq library construction

Tissue disaggregation was carried out adapting the protocol of Bertin et al [30]. Briefly, the frontoparietal bone, liver, heart and lung were dissected from euthanatized premetamorphic (NF58) tadpoles in PBS 0.6X. Each tissue was transferred to different tubes containing 0.6X HBSS (HyClone). The supernatants were discarded and replaced with 400 μL of 0.6X HBSS containing 0.1% collagenase P (Roche), and the tubes were gently agitated for 40 mins. The supernatants were discarded and replaced with 400 μL of 0.6X HBSS containing 0.1% collagenase P and 0.125% trypsin (Gibco), and the tubes were gently agitated for 10 mins. Finally, the supernatants were discarded and replaced with 400 μL of 0.6X HBSS containing 0.1% collagenase P and 0.125% trypsin, and the tubes were gently agitated for 15 mins. This last extraction step was performed four times, and all the supernatants were pooled in four tubes (a single tube per sample type) containing 1 mL of 0.6X L-15 (Leibovitz-15, HyClone). Cells were subsequently centrifuged at 500g for 7 mins at room temperature. After centrifugation, the supernatant was removed and the cells were resuspended in 50 μL of cold (4°C) 0.6X PBS and centrifuged at 500g for 5 mins at 4°C. Cells were resuspended in 50 µL of lysis buffer (10mM Tris-HCl, pH 7.4, 10mM NaCl, 3mM MgCl_2_, 0.1% Igepal), followed by pipetting up and down and incubating for 3 mins. A volume of 45 μL of lysed cells was diluted in 1 mL of PBS 0.6X and centrifugated at 500g for 10 mins at 4°C. During centrifugation, the remaining 5 μL of lysed cells was used to count the nuclei after trypan blue addition. After centrifugation, the supernatants were removed, and the nuclei were resuspended in 10 μL of PBS 0.6X. Approximately 80,000–100,000 nuclei per sample were transferred into fresh tubes and then centrifuged at 500g for 5 mins at 4°C. Again, the supernatants were carefully aspirated. Transposition reaction was performed following the Buenrostro et al. protocol [113]. The pelleted nuclei were gently resuspended with 50 µl of transposition reaction containing 25 µL 2X Tagmentation buffer (Illumina Cat FC-121-1030), 5 µL Tn5 transposase (Illumina Cat FC-121-1030) and 22,5 µL nuclease-free H2O and incubated at 37 °C for 30 mins. Transposed DNA was purified using a QIAGEN MinElute Reaction Cleanup kit (28204) and eluted with 10 μL of elution buffer (10 mM Tris-HCl, pH 8). The purified DNA was amplified using a reaction mixture containing 2X NEB PCR Master Mix (NEB M0541L), a universal forward primer i5, and an indexed reverse primer from the i7 Illumina series, with a PCR protocol of 72°C for 5 mins, 98°C for 30 s, and then eleven cycles of 98°C for 10 s, 63°C for 30 s,72°C for 1 min, with a final step at 72°C for 10 mins and hold at 4°C. PCR products were then cleaned up using AMPure beads (Agencourt APure XP, Beckman Coulter, A63880) to remove small DNA fragments according to the manual. ATAC-Seq libraries were then analyzed by Fragment Bioanalyzer and sequenced on an Illumina NextSeq 500 sequencer at the Transcriptomic and Genomic Marseille Luminy Platform (Aix-Marseille University, TAGC).

### Processing of ATAC-Seq data

Raw sequencing fastq files were assessed for quality, adapter content and duplication rates with FastQC v0.11.9 and trimmed using Cutadapt [114]. Trimmed reads were mapped against the *Xenopus tropicalis* reference genome (v9.1) using Bowtie2 [115] with the --very-sensitive parameter. Mapped reads whose quality was inferior to 40 and reads mapped to the mitochondrial genome were filtered out with SAMtools [116, 117]. Then, the “MarkDuplicates” function from Picard (Picard Tools, Broad Institute, 2018, https://broadinstitute.github.io/picard/) was used to remove duplicate reads and estimate library complexity. To obtain the NFR-associated reads, all reads larger than 120 bp were eliminated. Peak calling was performed using Genrich v0.6.1 (available at https://github.com/jsh58/Genrich“ \h) including two biological replicates per tissue with the parameters -j (ATAC-seq mode) -q 0.05 (FDR adjusted *p*-value). Peaks for the four tissues were merged into a standard peak list. The number of raw reads mapped to each standard peak was counted using the multicov function of BedTools v2.29.2 [118]. The edgeR package [119] was used to normalize the raw counts by TMM for each sample and the R function cor was used to calculate Pearson’s correlation between biological replicates. After the calculation of the scale factor, genome browser tracks in bigwig format were generated using deepTools bamCoverage [120].

### NFR and Mononucleosome arrangement

NucleoATAC with default parameters was applied to infer nucleosome positions for each tissue replicate [121]. TSS enrichment, plotting and clustering for NFR and mononucleosome were performed by the deepTools functions computeMatrix, plotProfile and plotHeatmap, respectively.

### Identification of tissue-specific open chromatin regions

To quantitatively measure the relative occupancy for each peak, we adopted a strategy based on Shannon Entropy to assign a tissue-specificity index to each element [10, 122, 123]. For each peak, we defined its relative accessibility in a tissue type i as Ri = Ei/ΣE, where Ei is the TMM value for the peak in the tissue i, ΣE is the sum of TMM values in all tissues. The entropy score for each peak across tissues can be defined as H = −1 * sum(Ri * log2Ri). An entropy score close to zero indicates the accessibility of this peak is highly tissue-specific. Based on the distribution of entropy scores, peaks displaying scores inferior to 1.2 were selected as tissue-restricted peaks.

### Tissue-specific and ubiquitous ATAC peaks analysis

HOMER v4.11 [124] was used to annotate genomic elements such as TSS and non-TSS (exonic, TTS, intergenic and intronic), determine the enrichment of transcription factor binding sites (TFBS), and calculate the %GC and %CpG from tissue-specific and ubiquitous peak sequences. Enriched Gene Ontology (GO) terms were found with topGO [125]. The *p*-value of GO analysis was adjusted with BH correction and only the top terms for tissue-specific and ubiquitous open chromatin regions were plotted.

To assess whether tissue-specific and ubiquitous peak sequences are conserved in other species, these regions were BLASTed against the following genomes using BLASTN (-outfmt 6, -task blastn, -max_target_seqs 1 parameters): *Homo sapiens* (hg19), *Xenopus laevis* (XenLae10.1), *Danio rerio* (denRa11), *Gallus gallus* (galGal6), *Callorhinchus milii* (calMil1), *Petromyzon marinus* (petMar3) and *Branchiostoma floridae* (braFlo1). BLASTN alignments inferior to 100 bp were removed and only those alignments with the best score were kept for each region of interest. Finally, peaks BLASTed with the hg19 genome were crossed with a list of annotated enhancers in nine different human cell lines (Gm12878, H1-hESC, HepG2, HMEC, HSMM, HUVEC, K562, NHEK, NHLF). Synteny determination and phenotyping analyzes were performed using GREAT v4 [126, 127].

### Sample collection and RNA-Seq library preparation

While liver tissue was disaggregated with a plastic pestle in TriReagent (Sigma), frontoparietal bone osteoblasts were extracted as described in Bertin et al [30] and then transferred to TriReagent. Homogenized samples were incubated for 5 mins at room temperature and centrifuged at 12,000g for 15 mins at 4°C. The supernatants were transferred to fresh tubes and 100 µL of BCP (1-bromo-3-chloropropane, Sigma) were added. Samples were incubated for 10 mins at room temperature and centrifuged at 12,000g for 15 mins at 4°C. The aqueous phase was transferred to a new Eppendorf tube and 125 µL of extraction buffer (1M Tris, 8M LiCl, 0.5M EDTA, 10% SDS) prewarmed at 56°C and 10 µL of proteinase K (10 mg/mL) were added and mixed. After 30 minutes of incubation, 250 µL of chloroform were added and mixed. The samples were incubated for 10 mins at 56°C and centrifuged at 12,000g for 15 mins at 4°C. The supernatants were transferred to fresh tubes and 250 µL of high salt precipitation solution (1.2 M NaCl, 0.8 M C_6_H_5_O_7_Na_3_) and 250 µL of isopropanol were added. The samples were mixed and incubated overnight at 20°C. Then, the samples were centrifuged at 12,000g for 10 mins at room temperature and the supernatants were discarded. The RNA pellets were washed with 300 µL of 70% ethanol, centrifuged at 12,000g for 5 mins at room temperature and finally dissolved in DEPC-treated water. The concentration and quality of the extracted RNA were determined using a NanoDrop Lite (Thermo). RNA-Seq libraries were constructed at the Transcriptomic and Genomic Marseille Luminy Platform (Aix-Marseille University, TAGC) using the TruSeq mRNA Library Prep Kit v2 (Illumina; California, USA). Libraries were paired-end sequenced on an Illumina NextSeq 500 sequencer.

### Processing of RNA-Seq data

Raw sequencing fastq files quality were assessed with FastQC, trimmed using Cutadapt and mapped against the *Xenopus tropicalis* transcriptome (v9.1) using HISAT2 [128]. Mapped reads whose quality was inferior to 40 and those reads mapped to the mitochondrial genome were filtered out with SAMtools [116, 117]. The number of raw reads mapped to each transcript was counted using the multicov function of BedTools v2.29.2 [118]. The edgeR package [119] was used to normalize the raw counts by TMM for both samples and R function cor to calculate Pearson’s correlation between biological replicates.

## Acknowledgements

This project was funded by a FONDECYT REGULAR 1190926 grant to SM, and by a FONDECYT REGULAR 1220708 and a FONDEQUIP EQM200056 to FA. We thank the Transcriptomics and Genomics Marseille-Luminy (TGML) platform for its excellent sequencing service. TGML is a member of the France Genomique consortium (ANR-10-INBS-0009). Work in the laboratory of SP was supported by recurrent funding from Institut National de la Santé et de la Recherche Médicale and Aix-Marseille University and Ligue contre le Cancer (Equipe Labellisée 2023). We are grateful to Hector Escriva and Stephanie Bertrand for their advice regarding the ATAC-Seq procedure.

## Supporting Information legends

**Supporting Information 1**: Fragment size distribution in linear and logarithmic (inset) scales for the first replicate of the liver **(A)**, heart **(B)**, and lung **(C)** samples. **(A’-C’)** For each tissue, the NFR (blue) and mononucleosome (red) signals at annotated *Xt* TSSs are visualized and separated into cluster 1 (left, clearly opened promoters harboring both high NFR and mononucleosome signals) and cluster 2 (right, closed promoters lacking a robust NFR signal and exhibiting a diffuse or absent mononucleosome signal).

**Supporting Information 2**: Genomic coordinates of the identified tissue-specific and ubiquitous nucleosome free regions (*Xenopus tropicalis* v9.1 genome).

**Supporting Information 3**: Comparison of the ATAC-Seq signals shown in 2kb windows for nucleosome-free regions (NFR) and mononucleosomes (Mono) at TSS **(A)** and non-TSS **(B)** tissue-specific and ubiquitous regions. For each type of region, the signal in bone, liver, heart, and lung is shown in red, brown, blue, and green, respectively.

**Supporting Information 4**: Frequency of TFBSs in 1kb windows centered on TSS **(A)** and non-TSS **(B)** ATAC-seq peaks. A color code indicates peaks that are specific to each tissue (bone, red; liver, brown, heart, blue) or ubiquitous (grey).

**Supporting Information 5:** Table summarizing a representative (non-exhaustive) bibliographical search for the known roles of transcription factors (TFs) binding to each of the enriched TFBSs identified at TSS and non-TSS peaks.

**Supporting Information 6:** Number of blasted TSS, non-TSS, and control regions (blasted sample size), as well as the corresponding number of obtained hits for each one of the analysed chordate genomes (*X.l.*, *Xenopus laevis*; *G.g.*, *Gallus gallus*; *H.s.*, *Homo sapiens*; *D.r.*, *Danio rerio*; *C.m.*, *Callorhinchus milii*; *P.m.*, *Petromyzon marinus*; *B.f.*, *Branchiostoma floridae*). For *H.s.*, we provide the total number of hits, as well as the number of hits that match a human region annotated as a transcriptional enhancer. N.d.: no data.

**Supporting Information 7**: Table showing the coordinates of the 46 *Xt* osteoblast-specific TSS that align to the TSS of their homologous gene in the human genome. The last column shows bibliographical references associating the corresponding genes to the osteogenesis process.

**Supplementary Information 8**: Genomic coordinates of the identified *Xenopus tropicalis* osteoblast-specific non-TSS NFRs conserved with the human genome.

**Supporting Information 9:** Comparison of TFBS enrichment between conserved and non-conserved osteoblast-specific non-TSS regions. **(A)** TFBS enrichment in 370 non-TSS osteoblast-specific *Xt* NFR conserved with humans. **(B)** TFBS enrichment at non-TSS osteoblast-specific NFR is compared between a set of *Xt* 370 regions conserved with humans (red), a set of *Xt* 6380 regions devoid of sequence conservation with human, chick, and elephant sharks (orange), and random control (grey).

**Supporting Information 10**: Result of GREAT v4 analysis showing the enrichment of human phenotypes, using the 370 *Xt* osteoblast-specific elements that are conserved with the human genome. Each phenotypic abnormality on the list is associated with mutations in the genes located closest to these elements.

**Supporting Information 11:** Table showing the coordinates of *Xt* osteoblast-specific non-TSS NFR conserved with humans and located at loci of genes associated to skeletal diseases according to the “Online Mendelian Inheritance in Man” database (OMIM, https://www.omim.org/).

**Supporting Information 12**: Bibliographical search for 15 genes whose TSS display a bone-specific ATAC-seq signal in *Xt* and that are evolutionarily conserved with human, chick and shark. Some representative, non-exhaustive, references reporting a function during osteogenesis and skeletogenesis are listed in the last column.

**Supporting Information 13**: Bibliographic search focused on osteoblastic development for 43 genes whose promoter is the most closely located to one of the 53 osteoblast-specific non-TSS ATAC-Seq peaks in *Xt* and that are evolutionarily conserved with annotated human enhancers, as well as with the chick and elephant shark genomes. Putative target genes were classified as transcription factors (“TF”, blue cells, 18 genes, 23 ATAC-Seq peaks), signaling pathways-related genes (“SIG”, green cells, 10 genes, 11 ATAC-Seq peaks), and as genes related to other cell processes (“OTH”, white cells, 15 genes, 19 ATAC-Seq peaks). Some representative, non-exhaustive, references reporting an osteoblastic function for the identified genes are listed in the last column.

**Supporting Information 14:** Representative, non-exhaustive, bibliographic search focused on stem cell function for genes closely located to *Xt* lung-specific non-TSS regions that are conserved with humans.

