## Supplementary material for "Architecture and evolutionary conservation of *Xenopus tropicalis* osteoblast-specific regulatory regions shed light on bone diseases and early skeletal evolution": Supp Data 1

**Supplementary Information 1**: Fragment size distribution in linear and logarithmic (inset) scales for the first replicate of the liver **(A)**, heart **(B)**, and lung **(C)** samples. **(A'-C')** For each tissue, the NFR (blue) and mononucleosome (red) signals at annotated *X.t.* TSSs are visualized and separated into cluster 1 (left, clearly opened promoters harboring both high NFR and mononucleosome signals) and cluster 2 (right, closed promoters lacking a robust NFR signal and exhibiting a diffuse or absent mononucleosome signal).

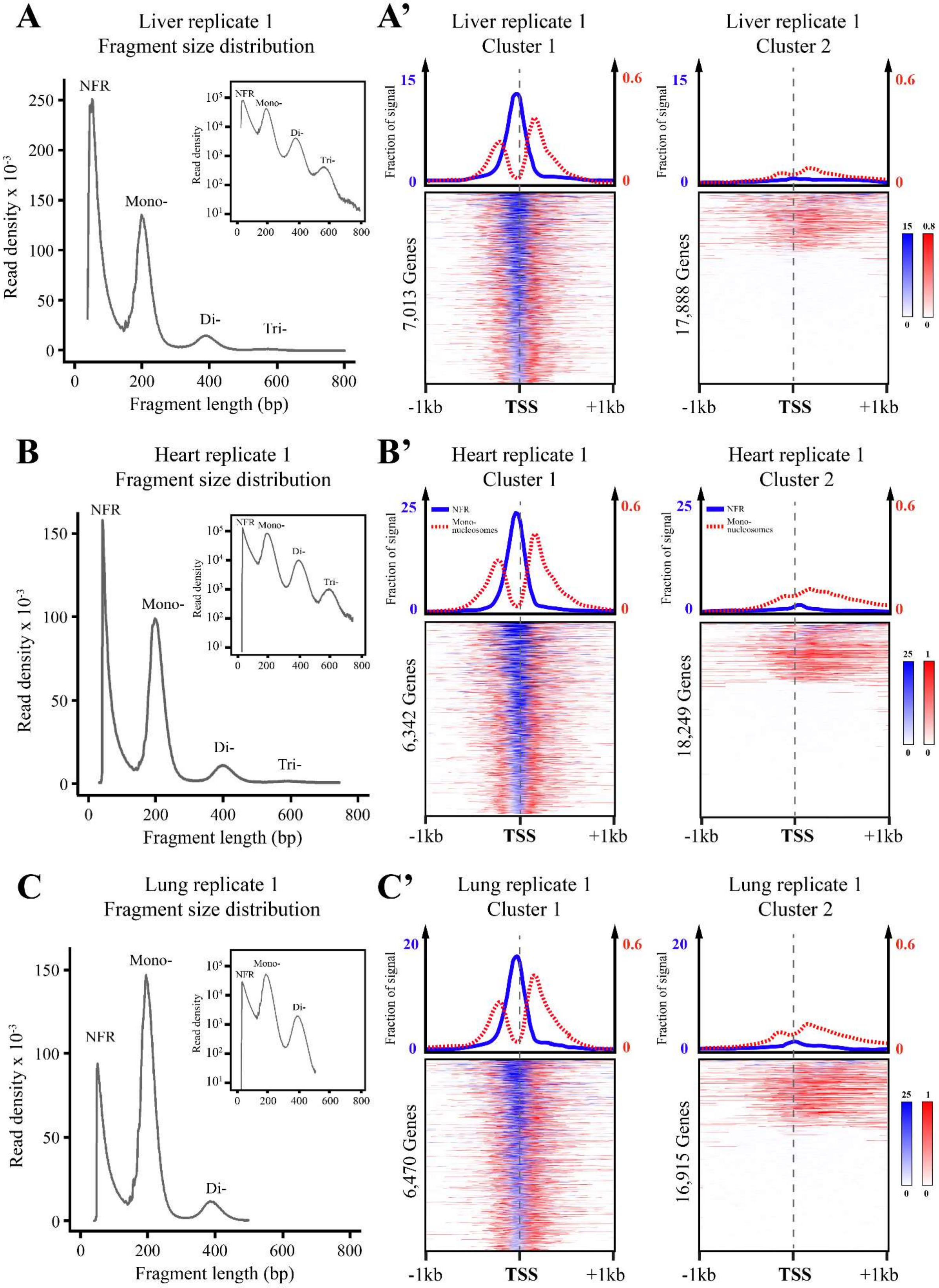
