## Supplementary material for "Architecture and evolutionary conservation of *Xenopus tropicalis* osteoblast-specific regulatory regions shed light on bone diseases and early skeletal evolution": Supp Data 3

**Supplementary Information 3**: Comparison of the ATAC-Seq signals shown in 2kb windows for nucleosome-free regions (NFR) and mononucleosomes (Mono) at TSS **(A)** and non-TSS **(B)** tissue-specific and ubiquitous regions. For each type of region, the signal in bone, liver, heart, and lung is shown in red, brown, blue, and green, respectively.

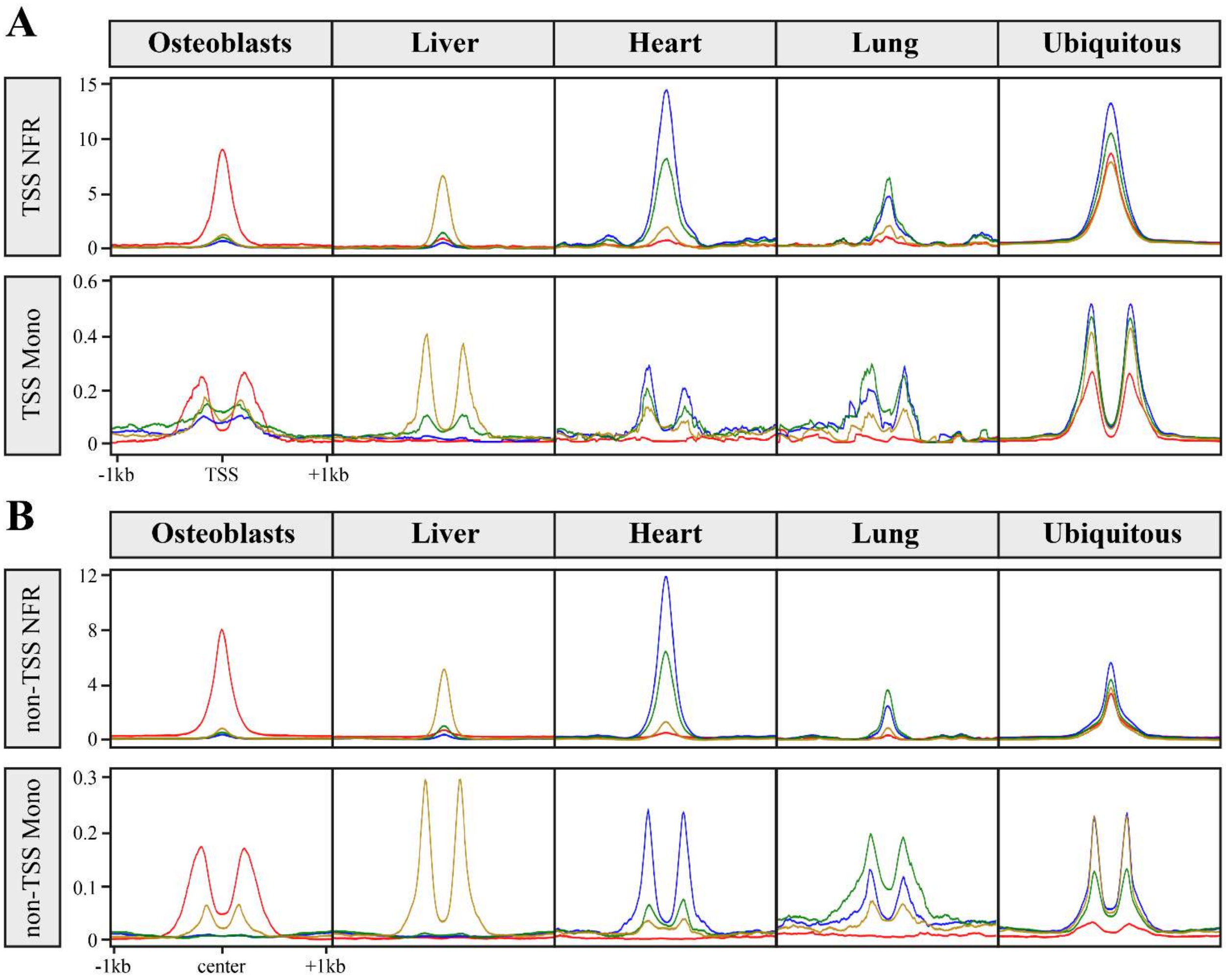
