## Supplementary material for "Architecture and evolutionary conservation of *Xenopus tropicalis* osteoblast-specific regulatory regions shed light on bone diseases and early skeletal evolution": Supp Data 4

**Supporting Information 4**: Frequency of TFBSs in 1kb windows centered on TSS **(A)** and non-TSS **(B)** ATAC-seq peaks. A color code indicates peaks that are specific to each tissue (bone, red; liver, brown, heart, blue) or ubiquitous (grey).

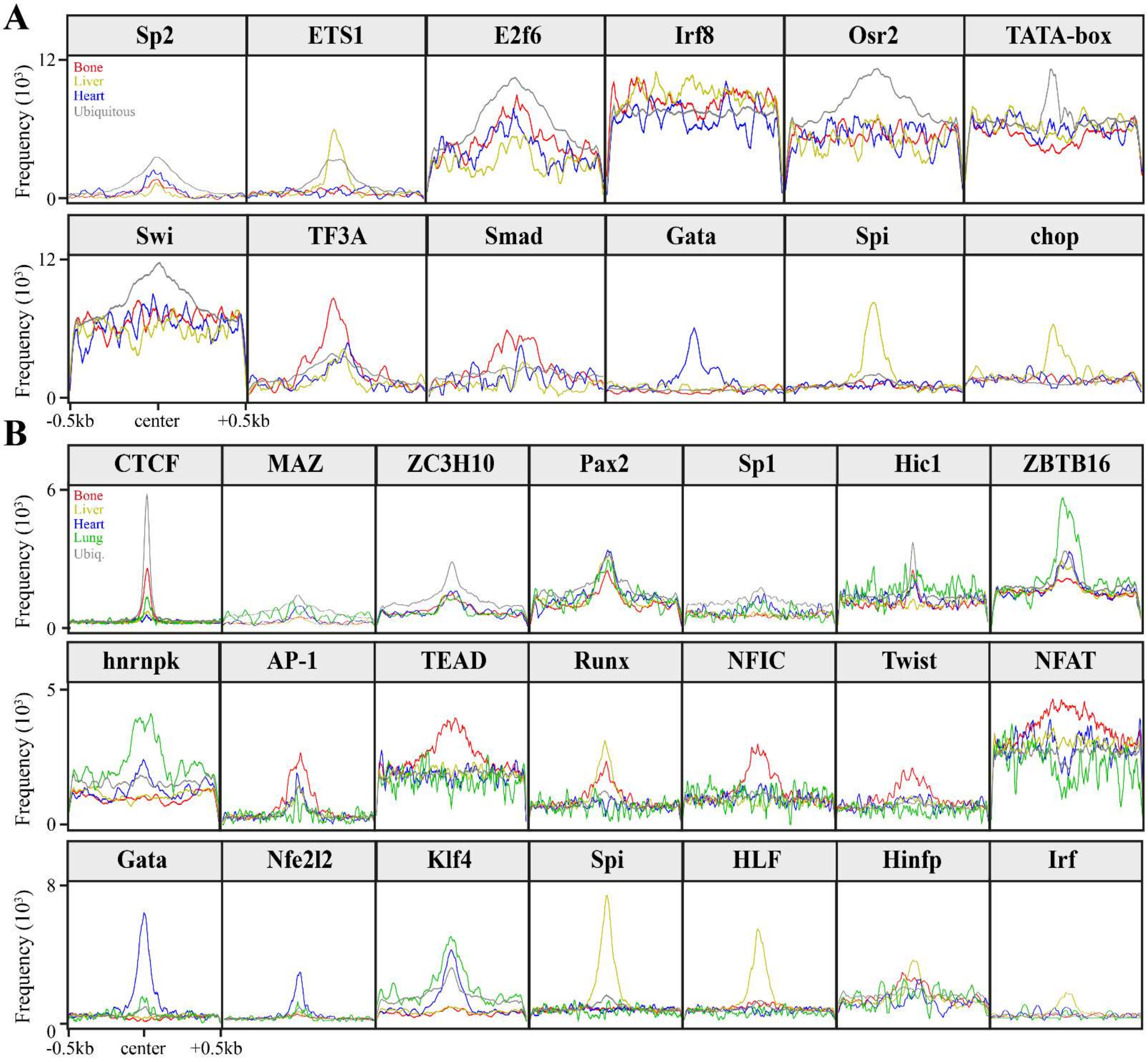
