## Supplementary material for "Architecture and evolutionary conservation of *Xenopus tropicalis* osteoblast-specific regulatory regions shed light on bone diseases and early skeletal evolution": Supp Data 5

**Supporting Information 5:** Table summarizing a representative (non-exhaustive) bibliographical search for the known roles of transcription factors (TFs) binding to each of the enriched TFBSs identified at TSS and non-TSS peaks.

<table>
<tr><td rowspan="3">Ubiquitous</td><td>TSS</td><td colspan="2">Sp2, E2f6 and TBP (TATA-box Binding Protein) function at ubiquitous and/or highly transcribed promoters [1-4].<br><br>By contrast, other enriched binding sites, including Meis [5, 6], Irf8 [7], Ets1 [8], Osr2 [9-11] and Swi [bound by Swi4 and Swi6 from the yeast HP1 family, see refs 12, 13, 14] are more difficult to interpret in the context of vertebrate ubiquitous promoter function.</td></tr>
<tr><td rowspan="2">non-TSS</td><td>Insulators and TAD architecture</td><td>CTCF and MAZ [15, 16].</td></tr>
<tr><td>Transcriptional enhancers</td><td>Zc3h10 as a metabolic regulator [17], Pax2/5/8 function at enhancers [18-20], binding of the ubiquitous Sp1 at enhancers [21], Hic1 function at enhancers [22].</td></tr>
<tr><td rowspan="2">Osteoblasts</td><td>TSS</td><td colspan="2">Smad: Smad1 as a BMP pathway effector in osteoblasts [23]. TFIIIA has so far only been reported to bind Pol III rRNA promoters [24].</td></tr>
<tr><td>non-TSS</td><td colspan="2">Osteoblastic role for the AP-1 member Fra1 [25], TEAD [26], Runx2 [27], NFIC [28], Twist1/2 [29] and NFAT [30].</td></tr>
<tr><td rowspan="2">Heart</td><td colspan="3">Cardiomyocytes: a role for Gata4 and Gata6 [31, 32].</td></tr>
<tr><td colspan="3">Erythrocytes: a role for Gata [33, 34], Klf [35, 36] and Nfe [33] transcription factors.</td></tr>
<tr><td>Liver</td><td colspan="3">TSS enrichment: one-third of CHOP binding events are located at promoters in a fibroblastoma cell line [37].<br>Role of SpiI, HLF, CHOP and Runx1 in macrophages and the myeloid lineage [38-45].</td></tr>
<tr><td>Lung</td><td colspan="3">Zbtb16 role in lymphoid lineage development [46].<br>hnRNP-K role in mouse embryonic stem cells [47].</td></tr>
</table>
