## Supplementary material for "Architecture and evolutionary conservation of *Xenopus tropicalis* osteoblast-specific regulatory regions shed light on bone diseases and early skeletal evolution": Supp Data 6

**Supporting Information 6:** Number of blasted TSS, non-TSS, and control regions (blasted sample size), as well as the corresponding number of obtained hits for each one of the analysed chordate genomes (*X.l.*, *Xenopus laevis*; *G.g.*, *Gallus gallus*; *H.s.*, *Homo sapiens*; *D.r.*, *Danio rerio*; *C.m.*, *Callorhinchus milii*; *P.m.*, *Petromyzon marinus*; *B.f.*, *Branchiostoma floridae*). For *H.s.*, we provide the total number of hits, as well as the number of hits that match a human region annotated as a transcriptional enhancer. N.d.: no data.

| | **Blasted sample size** | **Number of BLASTN hits against the genomes of the indicated chordate species** | | | | | | | | |
|---|---|---|---|---|---|---|---|---|---|---|
| | | ***X.l.* (L genome)** | ***X.l.* (S genome)** | ***G.g.*** | ***H.s.* (all)** | ***H.s.* (annotated enhancers only)** | ***D.r.*** | ***C.m.*** | ***P.m.*** | ***B.f.*** |
| **TSS peaks** | Osteoblasts-specific (524 regions) | 342 | 168 | 43 | 46 | N.d. | 15 | 24 | 6 | 5 |
| | Liver-specific (326 regions) | 205 | 88 | 7 | 8 | N.d. | 3 | 4 | 1 | 4 |
| | Heart-specific (218 regions) | 118 | 63 | 10 | 8 | N.d. | 5 | 7 | 7 | 3 |
| | Lung-specific (93 regions) | 46 | 20 | 6 | 6 | N.d. | 4 | 6 | 4 | 1 |
| | Ubiquitous (5945 regions) | 3847 | 1720 | 281 | 307 | N.d. | 208 | 179 | 136 | 79 |
| | Random (651 regions) | 283 | 196 | 21 | 12 | N.d. | 20 | 15 | 17 | 6 |

| | | | | | | | | | | |
|---|---|---|---|---|---|---|---|---|---|---|
| **Non-TSS peaks** | Osteoblasts-specific (6779 regions) | 4256 | 2094 | 417 | 370 | 127 | 113 | 173 | 42 | 24 |
| | Liver-specific (7259 regions) | 3495 | 1936 | 171 | 159 | 12 | 107 | 110 | 45 | 32 |
| | Heart-specific (2283 regions) | 1143 | 642 | 60 | 52 | 4 | 32 | 40 | 22 | 14 |
| | Lung-specific (804 regions) | 260 | 205 | 40 | 45 | 5 | 28 | 40 | 37 | 16 |
| | Ubiquitous (8706 regions) | 4434 | 2206 | 410 | 402 | 28 | 291 | 243 | 175 | 90 |
| | Random (6530 regions) | 2792 | 1848 | 134 | 114 | 8 | 158 | 163 | 186 | 60 |
