## Supplementary material for "Architecture and evolutionary conservation of *Xenopus tropicalis* osteoblast-specific regulatory regions shed light on bone diseases and early skeletal evolution": Supp Data 7

**Supplementary Information 7**: Table showing the coordinates of the 46 *X.t.* osteoblast-specific TSS that align to the TSS of their homologous gene in the human genome. The last column shows bibliographical references associating the corresponding genes to the osteogenesis process.

| ***Xenopus tropicalis*** **v9.1 genome coordinates** | | | ***Homo sapiens*** **hg19 genome coordinates** | | | | |
|---|---|---|---|---|---|---|---|
| **Chr** | **Start** | **End** | **Chr** | **Start** | **End** | **Gene Name** | **Known role in osteogenesis?** |
| Chr07 | 16.321.646 | 16.322.083 | chr10 | 128.076.961 | 128.077.098 | *adam12* | [1] |
| Chr01 | 106.482.909 | 106.483.454 | chr9 | 18.473.966 | 18.474.271 | *adamtsl1* | [2](Matrix organization) |
| Chr09 | 39.864.066 | 39.864.565 | chr2 | 189.839.018 | 189.839.234 | *col3a1* | [3] |
| Chr05 | 44.397.877 | 44.398.285 | chr6 | 132.272.497 | 132.272.720 | *ctgf* | [4, 5] |
| Chr02 | 102.210.169 | 102.210.447 | chrX | 13.835.102 | 13.835.292 | *gpm6b* | [6] |
| Chr06 | 32.929.051 | 32.929.295 | chr17 | 46.655.735 | 46.655.847 | *hoxa4* | [7] |
| Chr03 | 39.729.598 | 39.729.844 | chr12 | 102.874.254 | 102.874.500 | *igf1* | [8, 9] |
| Chr03 | 68.387.487 | 68.387.760 | chr7 | 102.553.295 | 102.553.421 | *lrrc17* | [10] |
| Chr05 | 56.774.298 | 56.774.815 | chr2 | 33.359.264 | 33.359.697 | *ltbp1* | [11, 12] |
| Chr08 | 90.867.330 | 90.867.791 | chr15 | 37.392.554 | 37.392.692 | *meis2* | [13] |
| Chr03 | 103.950.591 | 103.950.969 | chr15 | 96.873.778 | 96.873.986 | *nr2f2* | [14] |
| Chr01 | 177.081.485 | 177.081.770 | chr5 | 58.883.410 | 58.883.530 | *pde4d* | [15] |
| Chr06 | 75.463.633 | 75.463.947 | chr3 | 29.322.370 | 29.322.631 | *rbms3* | [16] |
| Chr05 | 143.433.745 | 143.434.128 | chr6 | 45.295.753 | 45.296.107 | *runx2* | [17-20] |
| Chr05 | 15.308.895 | 15.309.221 | chr2 | 45.236.597 | 45.236.788 | *six2* | [21] |
| Chr05 | 132.100.503 | 132.101.124 | chr2 | 5.832.854 | 5.833.044 | *sox11* | [22, 23] |
| Chr03 | 27.993.618 | 27.993.902 | chr5 | 141.704.578 | 141.704.706 | *spry4* | [24] |
| Chr05 | 118.067.405 | 118.067.747 | chr3 | 147.126.913 | 147.127.226 | *zic1* | [25, 26] |
| Chr06 | 95.220.837 | 95.221.549 | chr18 | 22.931.941 | 22.932.168 | *znf521* | [27-29] |
| Chr09 | 49.430.988 | 49.431.586 | chr2 | 204.193.221 | 204.193.360 | *abi2* | No |
| Chr01 | 149.189.551 | 149.189.795 | chr12 | 131.438.361 | 131.438.470 | *adgrd1* | No |
| Chr01 | 105.637.127 | 105.637.337 | chr9 | 16.705.027 | 16.705.152 | *bnc2* | No |
| Chr03 | 10.203.086 | 10.203.559 | chr7 | 134.464.011 | 134.464.352 | *cald1* | No |
| Chr03 | 16.281.156 | 16.281.408 | chr5 | 134.914.699 | 134.914.805 | *cxcl14* | No |
| Chr01 | 63.482.276 | 63.482.812 | chr4 | 105.416.241 | 105.416.340 | *cxxc4* | No |
| Chr06 | 122.929.460 | 122.929.847 | chr8 | 120.651.095 | 120.651.212 | *enpp2* | No |
| Chr02 | 18.097.555 | 18.097.841 | chr3 | 89.156.747 | 89.156.924 | *epha3* | No |
| Chr06 | 45.673.076 | 45.673.286 | chr7 | 14.029.239 | 14.029.421 | *etv1* | No |

| | | | | | | | |
|---|---|---|---|---|---|---|---|
| Chr08 | 66.182.222 | 66.182.497 | chr14 | 92.413.773 | 92.413.965 | *fbln5* | No |
| Chr04 | 12.222.749 | 12.223.259 | chr11 | 27.015.578 | 27.015.684 | *fibin* | No |
| Chr06 | 66.502.495 | 66.502.766 | chr7 | 41.740.441 | 41.740.626 | *inhba* | No |
| Chr02 | 130.591.941 | 130.592.214 | chr13 | 45.768.783 | 45.768.934 | *kctd4* | No |
| Chr08 | 41.244.078 | 41.244.334 | chrX | 86.772.592 | 86.772.697 | *klhl4* | No |
| Chr07 | 27.854.466 | 27.854.743 | chr10 | 95.517.517 | 95.517.645 | *lgi1* | No |
| Chr09 | 49.723.429 | 49.723.705 | chr3 | 185.080.946 | 185.081.042 | *map3k13* | No |
| Chr01 | 93.468.801 | 93.469.232 | chr4 | 87.281.095 | 87.281.329 | *mapk10* | No |
| Chr06 | 45.113.096 | 45.113.443 | chr7 | 15.726.219 | 15.726.408 | *meox2* | No |
| Chr01 | 33.329.466 | 33.329.959 | chr8 | 17.554.837 | 17.554.956 | *mtus1* | No |
| Chr01 | 52.729.970 | 52.730.576 | chr4 | 138.453.589 | 138.453.753 | *pcdh18* | No |
| Chr02 | 136.903.527 | 136.903.788 | chr12 | 57.736.154 | 57.736.255 | *r3hdm2* | No |
| Chr01 | 43.486.764 | 43.487.259 | chr4 | 160.188.808 | 160.188.949 | *rapgef2* | No |
| Chr08 | 81.267.417 | 81.267.804 | chr14 | 63.670.947 | 63.671.167 | *rhoj* | No |
| Chr05 | 24.416.052 | 24.416.345 | chr20 | 19.870.103 | 19.870.213 | *rin2* | No |
| Chr03 | 58.767.940 | 58.768.283 | chr7 | 116.660.146 | 116.660.308 | *st7* | No |
| Chr02 | 133.307.027 | 133.307.399 | chr12 | 49.582.754 | 49.582.913 | *tuba1b* | No |
| Chr02 | 117.356.784 | 117.357.942 | chr13 | 100.623.645 | 100.624.169 | *zic5* | No |
