## Supplementary material for "Architecture and evolutionary conservation of *Xenopus tropicalis* osteoblast-specific regulatory regions shed light on bone diseases and early skeletal evolution": Supp Data 9

**Supporting Information 9:** Comparison of TFBS enrichment between conserved and non-conserved osteoblast-specific non-TSS regions. **(A)** TFBS enrichment in 370 non-TSS osteoblast-specific *Xt* NFR conserved with humans. **(B)** TFBS enrichment at non-TSS osteoblast-specific NFR is compared between a set of *Xt* 370 regions conserved with humans (red), a set of *Xt* 6380 regions devoid of sequence conservation with human, chick, and elephant sharks (orange), and random control (grey).

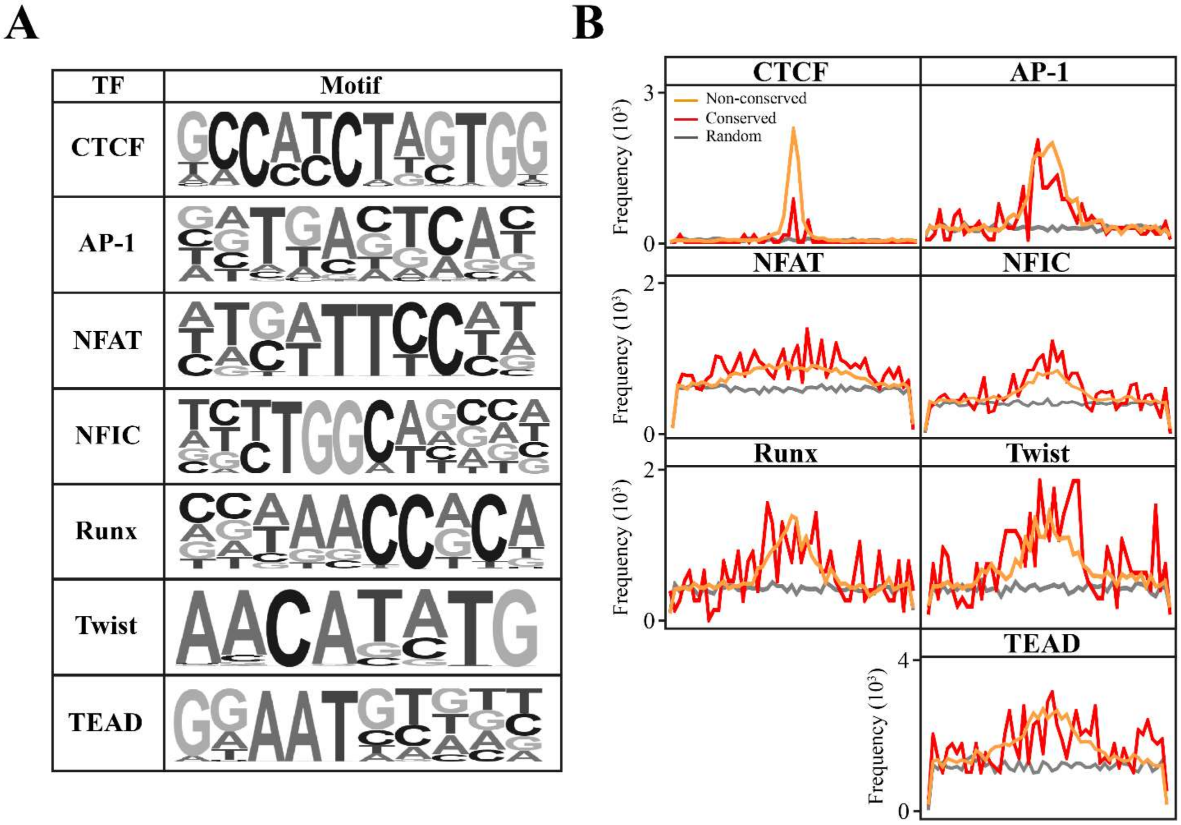
