## Supplementary material for "Architecture and evolutionary conservation of *Xenopus tropicalis* osteoblast-specific regulatory regions shed light on bone diseases and early skeletal evolution": Supp Data 10

**Supporting Information 10**: Result of GREAT v4 analysis showing the enrichment of human phenotypes, using the 370 *X.t.* osteoblast-specific elements that are conserved with the human genome. Each phenotypic abnormality on the list is associated with mutations in the genes located closest to these elements.

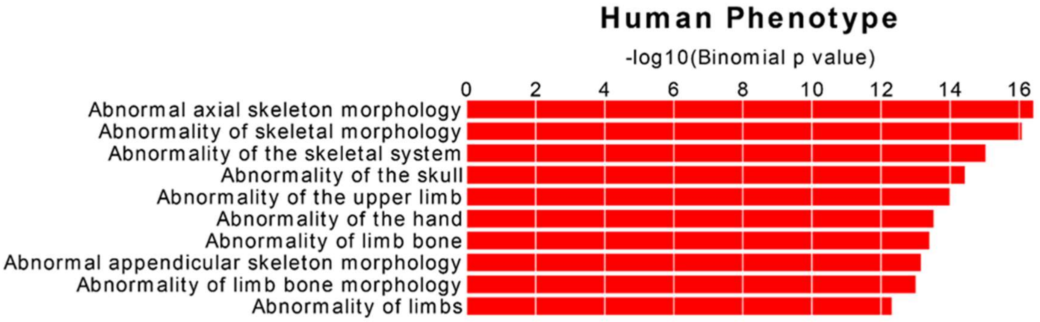
