## Supplementary material for "Architecture and evolutionary conservation of *Xenopus tropicalis* osteoblast-specific regulatory regions shed light on bone diseases and early skeletal evolution": Supp Data 11

**Supporting Information 11:** Table showing the coordinates of *Xt* osteoblast-specific non-TSS NFR conserved with humans and located at loci of genes associated to skeletal diseases according to the "Online Mendelian Inheritance in Man" database (OMIM, https://www.omim.org/).

| | | ***Xenopus tropicalis* v9.1 coordinates** | | | | **Human hg19 coordinates** | | | |
|---|---|---|---|---|---|---|---|---|---|
| **Locus / Putative target gene** | **Associated disease (OMIM number). Skeletal phenotype.** | **Chromosme / Scaffold** | **Start** | **End** | **Distance to putative target gene (bp)** | **Chromosme** | **Start** | **End** | **Distance to putative target gene (bp)** |
| ***flnb*** | Atelosteogenesis type III (108721). Skeletal dysplasia with vertebral fusions and craniofacial, carpal, tarsal, and phalangeal abnormalities. Disharmonious skeletal maturation, poorly modeled long bones, and joint dislocations. | scaffold_87 | 516.829 | 517.283 | 46.812 | chr3 | 58.088.076 | 58.087.927 | 93.875 |
| ***gorab*** | Geroderma osteodysplasticum (231070). Bowed long bones, and osteopenia with frequent fractures. | Chr04 | 108.081.635 | 108.082.226 | 44.150 | chr1 | 170.278.766 | 170.278.541 | 222.616 |
| ***gpc4*** | Keipert syndrome (301026). Craniofacial and digital abnormalities. | Chr08 | 46.817.871 | 46.818.110 | 8.989 | chrX | 132.546.362 | 132.546.588 | 3.043 |
| ***lemd3*** | Buschke-Ollendorff syndrome (166700). Affected individuals have osteopoikilosis consisting in osteosclerotic foci at the level of the epiphyses and metaphyses of long bones, wrist, foot, ankle, pelvis, and scapula. | Chr03 | 48.216.493 | 48.217.071 | 55.796 | chr12 | 65.721.501 | 65.721.079 | 157.728 |
| ***msx2*** | Craniosynostosis 2 (604757). Impaired skull growth, premature fusion of the cranial sutures. | Chr03 | 33.233.448 | 33.233.747 | 184.597 | chr5 | 173.811.879 | 173.811.639 | 339.777 |
| ***tcf12*** | Craniosynostosis 3 (615314). Impaired skull growth, premature fusion of the cranial sutures. | Chr03 | 90.017.727 | 90.018.199 | 18.219 | chr15 | 5.7545.981 | 5.7545.850 | 335.095 |
| ***trps1*** | Trichorhinophalangeal syndrome type I (190350). Craniofacial and skeletal abnormalities, including cone-shaped epiphyses at the phalanges, hip malformations, and short stature. | Chr06 | 120.820.455 | 120.820.898 | 83.960 | chr8 | 116.453.113 | 116.453.280 | 228.031 |
| | | | 120.841.339 | 120.841.635 | 63.150 | | 116.476.830 | 116.477.009 | 204.308 |
