## Supplementary material for "Architecture and evolutionary conservation of *Xenopus tropicalis* osteoblast-specific regulatory regions shed light on bone diseases and early skeletal evolution": Supp Data 12

**Supplementary Information 12**: Bibliographical search for 15 genes whose TSS display a bone-specific ATAC-seq signal in *X.t.* and that are evolutionarily conserved with human, chick and shark. Some representative, non-exhaustive, references reporting a function during osteogenesis and skeletogenesis are listed in the last column.

| ***Xenopus tropicalis* v9.1 genome coordinates** | | | ***Callorhinchus milii* calMil1 genome coordinates** | | | | |
|---|---|---|---|---|---|---|---|
| **Chromosome / Scaffold** | **Start** | **End** | **Scaffold name** | **Start** | **End** | **Gene Name** | **Known role in osteogenesis and skeletogenesis** |
| Chr08 | 66.182.223 | 66.182.497 | KI636070 | 507.881 | 508.009 | *fbln5* | *fbln5* in cartilage [1] and bone [2]. Also see [3] for a role of *fbln4* in bone. |
| Chr06 | 32.929.052 | 32.929.295 | KI635933 | 1.145.951 | 1.146.106 | *hoxa4* | Vertebral homeotic transformations [4-6]. Polycomb-dependent repression of Hox clusters during osteogenesis [7]. |
| Chr03 | 39.729.599 | 39.729.844 | KI635893 | 3.393.276 | 3.393.147 | *igf1* | *igf1* regulation in osteoblasts [8, 9]. |
| Chr01 | 93.468.802 | 93.469.232 | KI635904 | 661.633 | 661.470 | *mapk10* | Mitogen-Activated Protein Kinase Signalling in bone [10, 11]. |
| Chr04 | 97.340.947 | 97.341.340 | KI635895 | 3.660.090 | 3.660.385 | *mir199a* | *miR-199a* in osteoblasts and chondrocytes [12, 13]. |
| Chr05 | 143.433.746 | 143.434.128 | KI635960 | 1.225.578 | 1.225.318 | *runx2* | Initial mouse loss of function reports [14-17]. |
| Chr06 | 95.220.838 | 95.221.549 | KI635891 | 4.665.390 | 4.665.556 | *znf521* | *znf521* function in the osteoblastic lineage [18, 19]. |
| Chr01 | 43.486.765 | 43.487.259 | KI635921 | 2.802.817 | 2.802.943 | *rapgef2* | Not found |
| Chr03 | 10.203.087 | 10.203.559 | KI635973 | 1.415.880 | 1.415.697 | *cald1* | Not found |
| Chr01 | 174.898.792 | 174.899.277 | KI635918 | 1.585.495 | 1.585.618 | *nln* | Not found |
| Chr09 | 49.430.989 | 49.431.586 | KI635870 | 904.235 | 904.101 | *abi2* | Not found |
| Chr08 | 72.143.128 | 72.143.589 | KI635863 | 9.899.655 | 9.899.448 | *sav1* | Not found |
| scaffold_37 | 849.461 | 849.805 | AAVX02063905 | 189 | 326 | *aplp1* | Not found |
| Chr02 | 130.591.942 | 130.592.214 | KI635874 | 6.433.790 | 6.433.933 | *kctd4* | Not found |
| Chr02 | 117.356.785 | 117.357.942 | KI636011 | 1.225.881 | 1.225.562 | *zic5* | Not found |
