## Supplementary material for "Architecture and evolutionary conservation of *Xenopus tropicalis* osteoblast-specific regulatory regions shed light on bone diseases and early skeletal evolution": Supp Data 13

**Supplementary Information 13**: Bibliographic search focused on osteoblastic development for 43 genes whose promoter is the most closely located to one of the 53 osteoblast-specific non-TSS ATAC-Seq peaks in *X.t.* and that are evolutionarily conserved with annotated human enhancers, as well as with the chick and elephant shark genomes. Putative target genes were classified as transcription factors ("TF", blue cells, 18 genes, 23 ATAC-Seq peaks), signaling pathways-related genes ("SIG", green cells, 10 genes, 11 ATAC-Seq peaks), and as genes related to other cell processes ("OTH", white cells, 15 genes, 19 ATAC-Seq peaks). Some representative, non-exhaustive, references reporting an osteoblastic function for the identified genes are listed in the last column.

| ***Xenopus tropicalis* v9.1 genome coordinates** | | | ***Callorhinchus milii* calMil1 genome coordinates** | | | | | |
|---|---|---|---|---|---|---|---|---|
| **Chromosome / Scaffold** | **Start** | **End** | **Scaffold** | **Start** | **End** | **Putative target gene** | **Cellular Function** | **Known role in osteogenesis?** |
| Chr07 | 9.824.489 | 9.824.909 | KI635857 | 6.152.474 | 6.152.341 | *arid5b* | TF | No |
| Chr01 | 14.995.681 | 14.996.233 | KI635913 | 1.729.518 | 1.729.654 | *atoh8* | TF | [1] |
| Chr01 | 105.598.557 | 105.599.063 | KI635878 | 3.559.824 | 3.560.005 | *bnc2* | TF | [2, 3] |
| Chr06 | 32.078.043 | 32.078.382 | KI635933 | 473.069 | 473.243 | *creb5* | TF | No, but for cartilage see [4] |
| Chr07 | 13.997.643 | 13.997.936 | KI635857 | 1.392.764 | 1.392.559 | *ebf3* | TF | [5] |
| Chr04 | 129.993.970 | 129.994.312 | KI635892 | 4.615.013 | 4.614.789 | *foxp1* | TF | [6] |
| Chr03 | 57.629.539 | 57.629.933 | KI635889 | 1.105.170 | 1.105.470 | *foxp2* | TF | [7, 8] |
| Chr03 | 48.027.891 | 48.028.358 | KI635987 | 509.419 | 509.224 | *hmga2* | TF | [9, 10] |
| Chr03 | 48.196.698 | 48.197.077 | KI635987 | 399.252 | 399.099 | | | |
| Chr04 | 44.615.624 | 44.616.125 | KI635866 | 9.220.578 | 9.220.848 | *irx3* | TF | [11, 12] |
| Chr04 | 44.192.978 | 44.193.235 | KI635866 | 8.877.819 | 8.877.946 | | | |
| Chr01 | 170.373.368 | 170.373.594 | KI635881 | 315.718 | 315.872 | *mef2c* | TF | [13, 14] |
| Chr06 | 28.382.438 | 28.383.017 | KI635934 | 3.328.834 | 3.329.050 | *nfatc1* | TF | [15, 16] |
| Chr01 | 112.225.281 | 112.225.642 | KI635875 | 5.122.132 | 5.122.332 | *nfib* | TF | [17] |
| Chr01 | 160.771.065 | 160.771.767 | KI635923 | 1.385.093 | 1.384.688 | *prdm6* | TF | No, but for *prdm5* see [18] |
| Chr06 | 112.036.403 | 112.036.794 | KI635953 | 2.808.667 | 2.808.422 | *runx1t1* | TF | No |
| Chr05 | 75.880.264 | 75.880.523 | KI636156 | 333.945 | 334.164 | *sobp* | TF | No, but for craniofacial development see [19] |
| Chr04 | 22.546.403 | 22.547.098 | KI635866 | 6.130.392 | 6.130.975 | *tshz3* | TF | No, but for *tshz2* see [20] |
| Chr04 | 22.103.766 | 22.104.215 | KI635866 | 5.962.330 | 5.962.658 | | | |
| Chr09 | 73.238.671 | 73.238.907 | KI635900 | 3.407.929 | 3.407.772 | *zeb2* | TF | [21, 22] |
| Chr09 | 72.691.389 | 72.691.639 | KI635900 | 2.993.508 | 2.993.663 | | | |
| Chr06 | 107.438.012 | 107.438.257 | KI635861 | 3.404.604 | 3.404.744 | *zfhx4* | TF | [23] |
| Chr06 | 107.440.452 | 107.440.940 | KI635861 | 3.405.621 | 3.405.956 | | | |

| | | | | | | | | |
|---|---|---|---|---|---|---|---|---|
| Chr05 | 115.944.162 | 115.944.472 | KI635855 | 14.934.250 | 14.934.079 | *dock10* | SIG | No, but for the role of Rho GTPases see [24, 25] |
| Chr06 | 66.247.520 | 66.247.879 | KI635914 | 1.262.912 | 1.263.026 | *gli3* | SIG | For the role of the Hedgehog pathway see [26-29]. |
| Chr01 | 123.636.773 | 123.637.128 | KI635929 | 356.359 | 356.480 | *ptch1* | | |
| Chr01 | 123.434.834 | 123.435.137 | KI635929 | 161.557 | 161.684 | | | |
| Chr04 | 94.592.232 | 94.592.602 | KI635861 | 7.424.648 | 7.424.818 | *tgfbr3* | SIG | For the role of the BMP/Tgf-beta pathway see [30-32]. |
| Chr01 | 49.188.282 | 49.188.693 | KI635921 | 706.756 | 706.569 | *smad1* | | |
| Chr08 | 74.418.504 | 74.419.030 | KI635863 | 8.958.575 | 8.958.467 | *bmp4* | | |
| scaffold_628 | 49.644 | 49.843 | KI635860 | 7.735.534 | 7.735.652 | *gnas* | SIG | [33] |
| Chr01 | 33.129.119 | 33.129.451 | KI635879 | 5.692.910 | 5.693.056 | *mtmr7* | SIG | No |
| Chr01 | 56.178.409 | 56.178.835 | KI635950 | 3.165.111 | 3.165.252 | *spry1* | SIG | No, but for *spry2* see [34].<br>For the FGF pathway see [35, 36]. |
| Chr01 | 121.058.798 | 121.059.056 | KI635986 | 600.994 | 600.739 | *tle4* | SIG | [37] |
| Chr05 | 83.288.231 | 83.288.527 | KI635880 | 1.515.124 | 1.515.285 | *bach2* | OTH | No |
| Chr05 | 83.389.186 | 83.390.607 | KI635880 | 1.608.669 | 1.608.886 | | | |
| Chr02 | 88.445.800 | 88.446.310 | KI635901 | 3.232.523 | 3.232.314 | *bcas3* | OTH | No |
| Chr02 | 88.638.611 | 88.638.885 | KI635901 | 3.109.669 | 3.109.507 | | | |
| Chr03 | 58.438.485 | 58.438.831 | KI635889 | 1.812.886 | 1.813.010 | *cav2* | OTH | [38] |
| Chr06 | 23.679.875 | 23.680.192 | KI635858 | 14.447.600 | 14.447.463 | *cdk6* | OTH | [3, 39-41] |
| Chr01 | 9.169.690 | 9.170.028 | KI636023 | 999.314 | 999.155 | *dysf* | OTH | No |
| Chr06 | 121.482.976 | 121.483.297 | KI635909 | 4.279.169 | 4.279.413 | *eif3h* | OTH | No, but for eif2a see [42] |
| Chr09 | 66.007.981 | 66.008.663 | KI635868 | 923.863 | 924.183 | *fign* | OTH | No, but for fign-l1 see [43] |
| Chr02 | 146.185.540 | 146.185.875 | KI635874 | 133.350 | 133.585 | *fndc3a* | OTH | No |
| Chr05 | 24.453.800 | 24.454.251 | KI635915 | 3.285.229 | 3.285.359 | *naa20* | OTH | No |
| Chr02 | 27.454.082 | 27.454.681 | KI635901 | 2.097.142 | 2.097.006 | *smg6* | OTH | [44, 45] |
| Chr02 | 27.476.410 | 27.476.810 | KI635901 | 2.068.440 | 2.068.309 | | | |
| Chr02 | 27.545.358 | 27.545.572 | KI635901 | 2.036.755 | 2.036.579 | | | |
| Chr05 | 37.440.947 | 37.441.775 | KI635888 | 2.004.956 | 2.005.158 | *syne1* | OTH | No |
| Chr05 | 59.276.176 | 59.277.060 | KI635885 | 297.927 | 298.300 | *tomm20* | OTH | No |
| Chr05 | 135.090.827 | 135.091.439 | KI635905 | 3.926.593 | 3.926.257 | *lpin1* | OTH | No |
| Chr06 | 101.405.540 | 101.405.778 | KI635902 | 408.551 | 408.742 | *ubxn2b* | OTH | No |
| Chr01 | 105.270.809 | 105.271.074 | KI635878 | 3.289.937 | 3.290.060 | *ccdc171* | OTH | No |
