## Supplementary material for "Architecture and evolutionary conservation of *Xenopus tropicalis* osteoblast-specific regulatory regions shed light on bone diseases and early skeletal evolution": Supp Data 14

**Supporting Information 14:** Representative, non-exhaustive, bibliographic search focused on stem cell function for genes closely located to *X.t.* lung-specific non-TSS regions that are conserved with humans.

| Chromosome / Scaffold | Start | End | Annotation | Distance to nearest TSS (pb) | Gene Name | Reported role in stem cells | Refs. |
|---|---|---|---|---|---|---|---|
| Chr10 | 5.100.511 | 5.100.750 | exon | 934 | *hoxb8* | YES, IN LUNGS | [1] |
| Chr04 | 61.064.962 | 61.065.163 | exon | 17.569 | *cdh15* | YES | [2] |
| Chr09 | 62.326.721 | 62.327.078 | exon | 742 | *dlx1* | YES | [3] |
| Chr02 | 81.596.703 | 81.596.938 | Intergenic | 8.627 | *lhx3* | YES | [4] |
| scaffold_39 | 388.518 | 389.425 | exon | 165 | *msl1* | YES | [5, 6] |
| Chr05 | 38.615.443 | 38.615.797 | TTS | 225 | *ovol2* | YES | [7] |
| Chr03 | 108.739.938 | 108.740.200 | TTS | 347 | *purb* | YES | [8] |
| Chr08 | 34.184.933 | 34.185.379 | exon | 807 | *sox3* | YES | [9] |
| Chr10 | 22.446.850 | 22.447.124 | TTS | 624 | *sox9* | YES | |
| scaffold_418 | 39.415 | 39.665 | exon | 3.432 | *tcfl5* | YES | [10] |
| Chr03 | 31.163.042 | 31.163.836 | exon | 112 | *adrb2* | NO | |
| Chr02 | 95.138.287 | 95.138.526 | intron | 36.248 | *auts2* | NO | |
| Chr09 | 32.606.771 | 32.607.318 | Intergenic | 10.407 | *ercc4* | NO | |
| Chr01 | 17.005.725 | 17.005.957 | intron | 18.567 | *fbxl5* | NO | |
| scaffold_555 | 13.126 | 13.555 | exon | 12.513 | *fbxo42* | NO | |
| Chr10 | 7.891.392 | 7.891.578 | intron | 9.756 | *kcnh2* | NO | |
| Chr04 | 15.878.384 | 15.878.686 | TTS | 395 | *kcnq1* | NO | |
| Chr05 | 11.110.536 | 11.110.938 | exon | 7.857 | *nek2* | NO | |
| Chr01 | 79.507.797 | 79.508.028 | exon | 791 | *onecut3* | NO | |
| Chr10 | 3.826.106 | 3.827.062 | exon | 448 | *pip4k2a* | NO | |
| scaffold_788 | 17.818 | 18.015 | intron | 3.769 | *pla2g2d* | NO | |
| Chr09 | 8.865.949 | 8.866.148 | intron | 7.216 | *sec14l1* | NO | |
| scaffold_4201 | 4.970 | 5.199 | TTS | 1.582 | *slc15a5* | NO | |
| Chr05 | 22.578.803 | 22.579.020 | Intergenic | 4.692 | *snap25* | NO | |
| scaffold_6690 | 3.555 | 3.726 | TTS | 2.286 | *tpd52l2* | NO | |
| Chr05 | 39.521.918 | 39.522.211 | exon | 1.150 | *trim67* | NO | |
| Chr01 | 79.823.057 | 79.823.249 | exon | 27.888 | *unc13a* | NO | |
| Chr10 | 37.782.908 | 37.783.122 | exon | 273 | *uts2r* | NO | |
| Chr05 | 69.635.875 | 69.636.416 | intron | 1.705 | *vgll2* | NO | |
| Chr09 | 73.422.108 | 73.422.379 | intron | 10.535 | *zeb2* | NO | |
